# Niche theory for within-host parasite dynamics: Analogies to food web modules via feedback loops

**DOI:** 10.1101/2021.08.15.456318

**Authors:** Ashwini Ramesh, Spencer R Hall

## Abstract

Why do parasites exhibit a wide dynamical range within their hosts? For instance, why can a parasite only sometimes successfully infect its host? Why do some parasites exhibit large fluctuations? Why do two parasites coinfect, exclude each other, or win only sometimes over another (via priority effects)? For insights, we turn to food webs. An omnivory model (IGP) blueprints one parasite competing with immune cells for host energy (PIE), and a competition model (keystone predation, KP) mirrors a new coinfection model (2PIE). We then draw analogies between models using feedback loops. We translate those loops into the intraspecific direct (DE) and indirect effects (IE) that create various dynamics. Three points arise. First, a prey or parasite can flip between stable and oscillatory coexistence with their enemy with weakening IE and strengthening DE. Second, even with comparable loop structure, a parasite cannot exhibit priority effects seen in IGP due to constraints imposed by production of immune cells. Third, despite simpler loop structure, KP predicts parallel outcomes in the two-parasite model due to comparable structure of interactions between competing victims and their resources and enemies. Hence, food web models offer powerful if imperfect analogies to feedbacks underlying the dynamical repertoire of parasites within hosts.

## INTRODUCTION

Parasites can show a variety of dynamics within hosts. These within-host dynamics can significantly affect health of individual hosts (Råberg *et al* 2009; Cressler *et al* 2014; Bashey 2015) and alter population-level disease outbreaks (Mideo 2008; Råberg *et al* 2012; Gorisch *et al* 2018). For instance, exposure to parasite propagules often causes infection of some but not all hosts despite seemingly similar starting conditions (Merrill & Cáceres 2018). The success or failure of invasion and persistence of a parasite within a host determines its infection status or even death (Duneau *et al* 2017; Merrill & Cáceres 2018). Once infected, the stability or instability of within-host dynamics can further impact both host health and transmission to others. For instance, periods of high abundance of a cycling gut parasite can both decrease foraging and increase transmission to bee hosts (Otterstatter & Thomson 2006). Similarly, coinfection (*i.e.,* within-host coexistence of parasites) can reduce fecundity of hosts more than single infection (Lohr *et al* 2010). At the between-host scale, coinfection can reduce or increase outbreaks compared to singly infected populations (Clay *et al* 2019; Susi *et al* 2015). Consequently, various dynamics of parasites can arise within hosts, likely mediated by immune systems and nutrient/energy availability (Graham 2008; Cressler *et al* 2014). But which mechanisms determine those dynamics?

To answer this question, two classic food web modules from community ecology provide analogies. In intraguild predation (IGP), an omnivorous predator and its prey compete for a shared resource (Holt & Polis 1997; Verdy & Amarasekare 2010). Similarly, immune cells and a parasite compete for shared energy within a host (Fig. 1A - C; Cressler *et al* 2014). Hence, mechanisms that capture the repertoire of dynamics in IGP (stable coexistence, oscillations, priority effects, exclusion) could also produce those observed in single-parasite studies (Otterstatter & Thomson 2006; Merrill & Cáceres 2018). Additionally, in the diamond-keystone predation (KP) model, two prey species share a resource (exploitative competition) and a predator (apparent competition: Holt *et al*. 1994, Leibold 1996). Similarly, two parasites can compete for a shared resource while facing immune attack (Fig. 1D - F). In such a scenario, the KP model could anticipate the tradeoffs and niche dimensions needed for coexistence, exclusions, and priority effects of coinfecting parasites (Natsopoulou *et al* 2015). The underlying hope, then, is that food web modules might provide apt blueprints, because they share basic consumer-resource structure with their within-host analogues.

**Figure 1:**
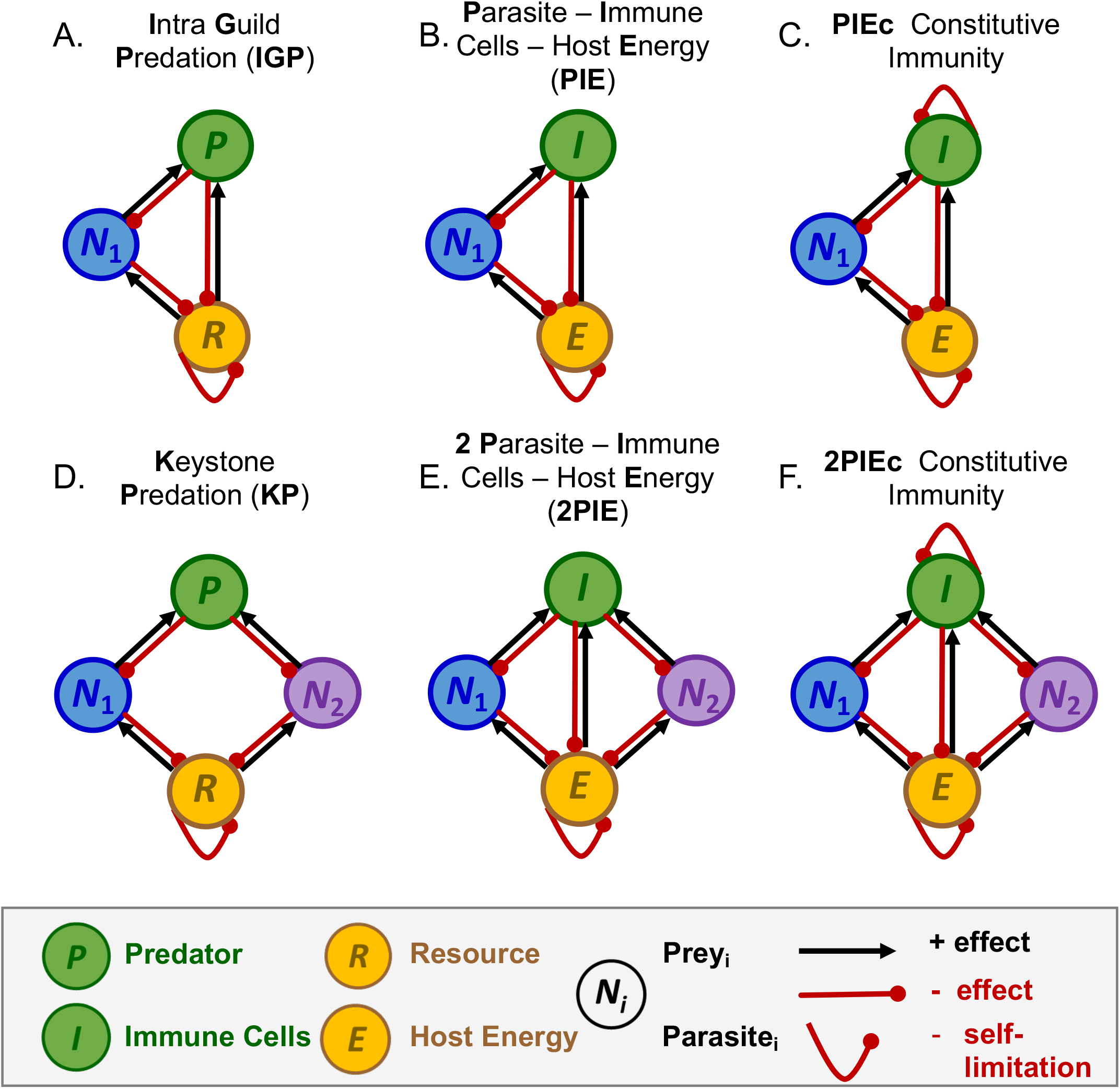
Direct interspecific effects and self-limitation in food web and within-host parasite models. Top row, single prey or parasite models (three-dimension, 3D): (A) one prey-predator-resource (intraguild predation, IGP), (B) one parasite-immune cells-host energy (PIE), and (C) PIEc with constitutive immunity (i.e., fixed allocation of energy to production of immune cells). *Bottom row, two prey or two parasite models (four dimension, 4D):* (D) two prey - predator-resource (keystone predation, KP), (E) two parasite - immune cells - host energy (2PIE), and (F) 2PIEc with constitutive immunity. All direct effects evaluated when densities are positive (i.e., at a feasible interior equilibrium). Red (black) arrows: negative (positive) interspecific direct effect; red curve: negative intraspecific specific effect.

However, a major complication arises: the enemy is generated differently in food webs vs. within a host (Wodarz 2006). Predators attack and assimilate prey for reproduction into new predators. Immune cells also attack parasites, but they simultaneously require host energy to produce immune cells (Cressler *et al* 2014). These “consumer-resource like” interactions can involve products of immune cells, parasites, and energy. Additionally, hosts can allocate a baseline level of energy straight to immune cells, a pipeline not enjoyed by even omnivorous predators in typical food webs. Therefore, these fundamentals of immune function – even when highly simplified - create more interaction links (Fig. 1). Those additional links might alter feedbacks underpinning the range of stability outcomes (Metcalf *et al* 2020), possibly undermining analogies from food webs or creating new outcomes altogether. Given these issues (Wodarz 2006; Alizon *et al* 2008; Fenton & Perkin 2010), how could we usefully compare and contrast stability outcomes in food webs (IGP, KP) to their within-host analogues?

We tackle this challenge by using a feedback loop approach. Feedback loops link the strength of consumer-resource interactions to stability (Puccia & Levins 1991). Indeed, feedback loops characterize the biology behind stability and enable comparison of structurally similar but biologically distinct systems. Every *n-*dimensional system consists of *n* levels of feedback (*F_n_*). The simplest level of feedback, level 1 (*F_1_*), involves intraspecific direct effects (DE). Here, a small increase of a species impacts its own growth rate – positively or negatively - via interaction with itself and from those directly consuming it. Then, longer loops (from 2 through *n* dimensions) involve combinations of intra- and inter-specific direct effects (calculated from Jacobian matrices). For instance, in a two-species loop, an increase in a resource’s density could increase growth rate of its consumer. Increased density of its consumer, in turn, could then depress growth rate of that resource (hence, binary consumer-resource interactions create negative *F_2_*). With more dimensions, the connections between interactors become more complex and varied, potentially introducing both positive and negative feedbacks. The sign and strength of feedback at all relevant levels, then, governs stability. Additionally, key ratios of those longer loops determine intraspecific indirect effects (IE; developed in Box 1). As we then show, feedback at certain levels, translated into intraspecific direct and indirect effects, determines stability of the interactions.

Using intraspecific direct and indirect feedback loops, we first revisit the foundational IGP and KP models. We then apply these loop-based lessons to analogous parasite-immune-energy models (PIE and 2PIE; Fig. 1). To study these analogues, we assume that parasite(s) respond to attack by a common immune system. That assumption surely oversimplifies some examples (*e.g.,* in vertebrates: Ezenwa & Jolles 2011), but it seems like a reasonable place to start for others (Graham 2008; Griffits *et al* 2015; Rynkiewicz *et al* 2015). Additionally, we assume that immune cells and parasites share host energy (a resource, following Griffits *et al* 2015). We then evaluate how host energy and immune system interact to mediate parasite dynamics within hosts (De Roode *et al* 2005; Ezenwa & Jolles 2011; Doublet *et al* 2015).

The analysis involves three parts. First, we translate loops into a scheme framed as intraspecific direct and indirect effects (DE and IE). This interpretation works most thoroughly for 3D systems (IGP and PIE models), but we extended key ideas to 4D systems (KP and 2PIE models). Second, using these tools, we analyzed stability outcomes for IGP vs. PIE. We find that IGP and PIE share some dynamics (coexistence, stably or via oscillations) due to similar loop structure. However, the nature of the generation of immune cells prevents some outcomes present in IGP (involving two variations on alternate stable states). Third, we find that despite having simpler feedback links, KP anticipates coexistence/coinfection vs. priority effects (or alternate stable states) of competing parasites in 2PIE models. Furthermore, these outcomes arise for conceptually similar reasons. Specifically, depending on their traits, competing prey or parasites exhibit symmetries (coexistence/coinfection) or asymmetries (priority effects) in ratios describing ‘*effects on’* their enemies and resources vs. how they are ‘*affected by*’ them. Those (a)symmetries determine the strength of tradeoffs and competitive hierarchies among competitors. They also underpin the sign of feedback at a key level, hence they govern whether parasites experience negative or positive indirect effects. Furthermore, allocation of energy to constitutive immunity enhances negative feedback, thereby shrinking regions of within-host oscillations, coinfection, and parasite burden. Thus, feedback loops, guided by our interpretation schemes, show when and why comparable dynamics arise in these food webs and their within-host analogues. However, we highlight why differences in production of enemies constrain some outcomes within hosts.

## METHODS AND RESULTS

### Overview of the models

We compare and contrast modules of food web and within-host dynamics. The model with one prey, predator, and resources (intraguild predation, IGP; Holt & Polis 1997; Verdy & Amarasekare 2010) parallels a within-host model of one <u>p</u>arasite species, immune cells, and host <u>e</u>nergy (PIE; modified from Hite & Cressler 2018). Similar, a two prey, predator, resource model (keystone predation, KP; Leibold 1996) parallels one with the new two parasite species, immune cells, and host energy model (2PIE). However, while the food web and within-host modules share similar structure, they create enemies differently. Specifically, predators proliferate via direct consumption of their prey, while immune cells jointly require both parasites and a shared energy. Additionally, the within-host modules include two variants that commonly arise in invertebrate host-parasite systems. In one, only parasites activate production of immune cells (PIE and 2PIE models). In the other, energy is continuously allocated to maintain baseline immune function, even without parasites (PIEc and 2PIEc, constitutive immunity). Despite their differences, we compare across food web and within-host modules using feedback loops. Feedback loops provide a common metric to unpack the biology underlying stability. Jacobian matrices provide the starting point (Box 1, Appendix Section 1). Each of Jacobian terms represents the direct effect of species *j* on growth rate of species *i* (*J_ij_*), yielding (hereafter) interspecific *positive effects* (black arrow) and *negative effects* (red arrow) and intraspecific *self-limitation* (red curve) or *self-facilitation* (black curve; Box 1, Fig. 1 [not seen here]). As shown below, these terms combine into loops at various levels.

### One prey - predator - resource (IGP) | One parasite - immune cells - host energy (PIE & PIEc)

At their heart, both IGP and PIE models hinge on binary consumer-resource like interactions (Fig.2, Table 1). In these interactions, consumers directly benefit while resources suffer direct costs. In IGP, omnivorous predators (*P)* can consume prey (*N_1_*) and resources (*R*). An interior equilibrium becomes feasible when each species can maintain positive density (*P\** > 0, *N*_1_*\** > 0, *R\** > 0). At IGP’s interior, these consumer-resource interactions exert positive effects on the predator but negative ones on prey and resource. Then, prey also indirectly compete with predators for the shared resource. Consumption of resources benefits the prey and harms resources. Finally, chemostat-like renewal imposes self-limitation on resources (Fig. 1A). Similarly, in the within-host models, immune cells (*I*) “consume” two items, simultaneously killing parasites (*N*_1_) while taking up host energy (*E*). At PIE’s interior, these consumer-resource-like interactions have a positive effect on *I* and negative on *N*_1_ and *E*. The parasite competes for this shared energy; consumption of host energy positively effects parasites and negatively effects energy. Additionally, the donor-controlled renewal of host energy imposes self-limitation on energy without baseline allocation (*a_b_* = 0; Fig. 1B). Fixed energy allocated to immune cells (*a*_b_ > 0) creates additional self-limitation on the immune cells in PIEc (Fig. 1C).

**Figure 2:**
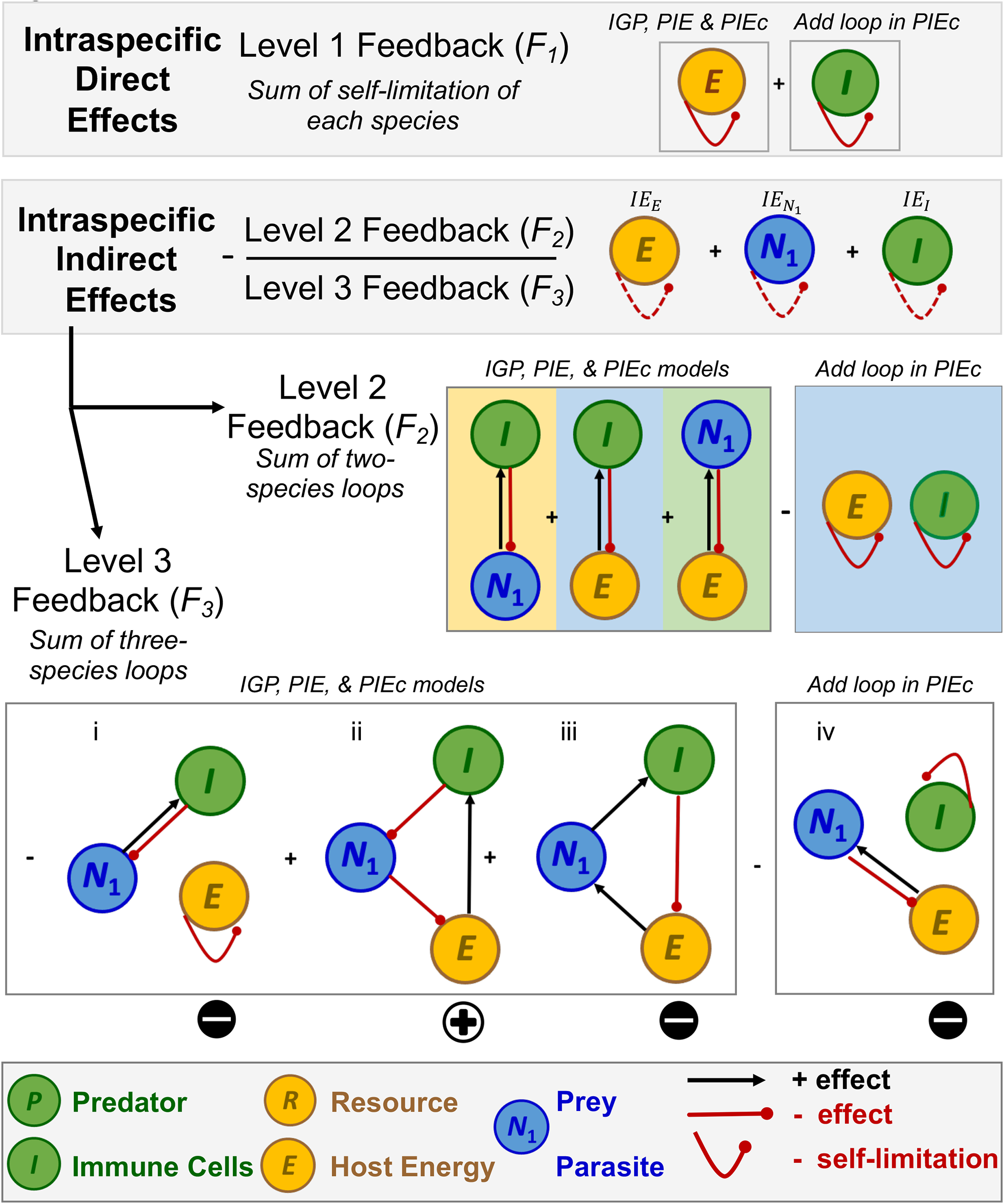
Intraspecific direct and indirect effects: an illustration of levels of feedback in IGP, PIE, and PIEc models (Fig. 1A-C). Evaluated at a feasible interior equilibrium, the IGP and PIE models have similar feedback structure. PIEc adds additional loops involving immune self-limitation. Level 1 feedback (*F_1_*) is the sum of the shorter intraspecific direct effects (solid curves): self-limitation (-) of resource/energy and immune cells (PIEc only). In intraspecific indirect effects (dashed curves), each species limits or facilitates itself via interaction with other species in longer loops. The intraspecific indirect effect (IE) sums that of energy, *E* (IE_E_; yellow shading), parasite *N_1_* (IE_N_1__; blue), and immune cells *I* (IE_I_; green). IE, in turn, is a ratio of two levels of feedback. Level 2 feedback (*F_2_*, numerator of IE) sums pairwise consumer-resource loops (-) and the product of energy and immune self-limitation (-; only in PIEc; these are the numerators of the IE components). Level 3 feedback (*F*_3_, denominator of IE) sums three-species loops. These loops from *L-R* are: (i) “*N_1_ is eaten*”, the *I-N_1_* loop times *E* self-limitation (-), (ii) “*starving the enemy*” (+), (iii) “*feeding the enemy*” (-), and (iv) “*N_1_ eats*”, the *E-N_1_* loop times *I* self-limitation (-; PIEc only). This sum of (i)-(iv) determines the sign of *F_3_* and of IE (since IE = *–F_2_*/*F_3_*, and *F_2_* < 0 always). The predator (*P*, green), prey (*N_1_*, blue), and resources (*R*, yellow) in IGP are analogous to immune cells (*I*), parasite (*N_1_*), and host energy (*E*), respectively, in the PIE models.

**Table 1.**
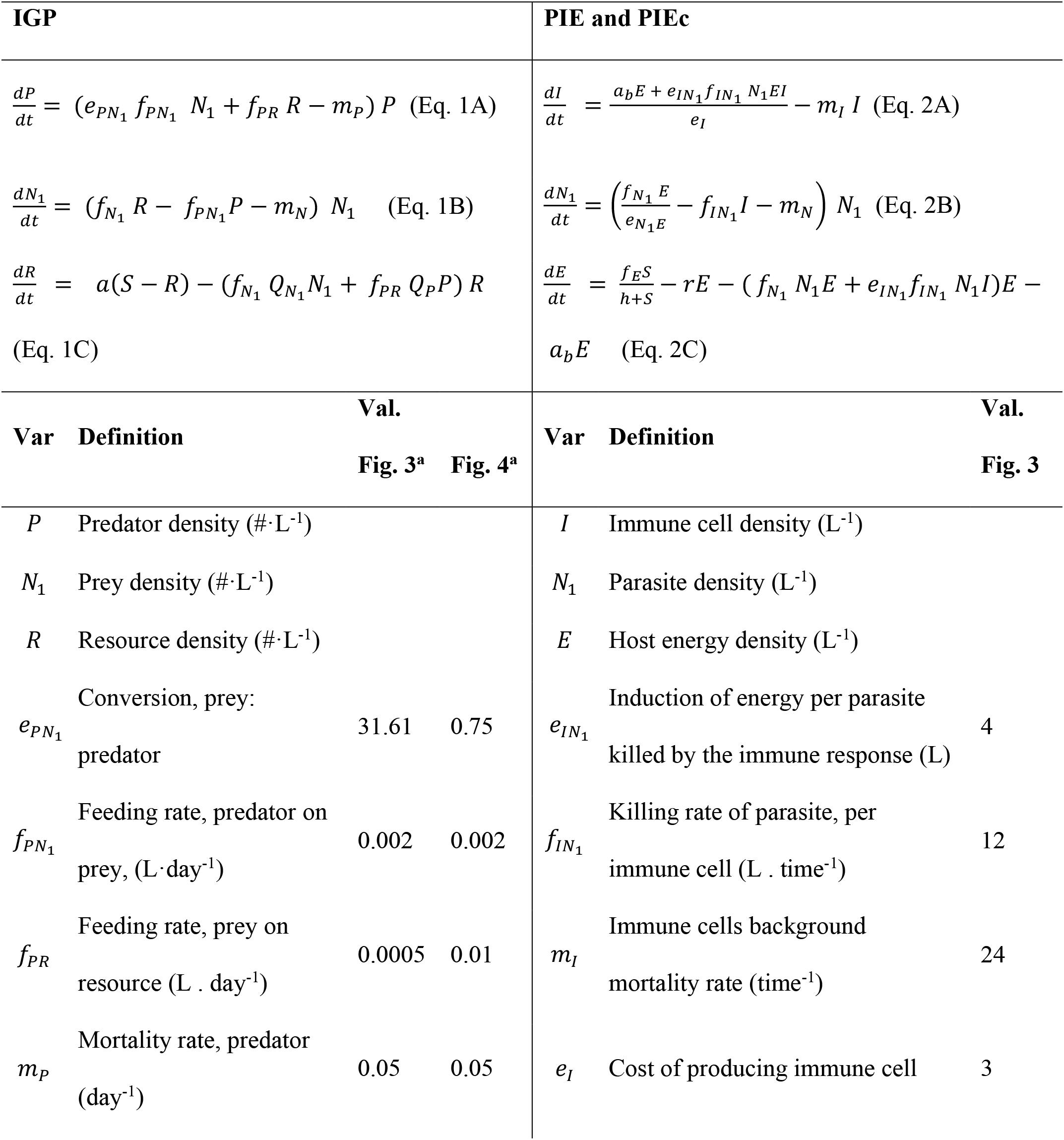
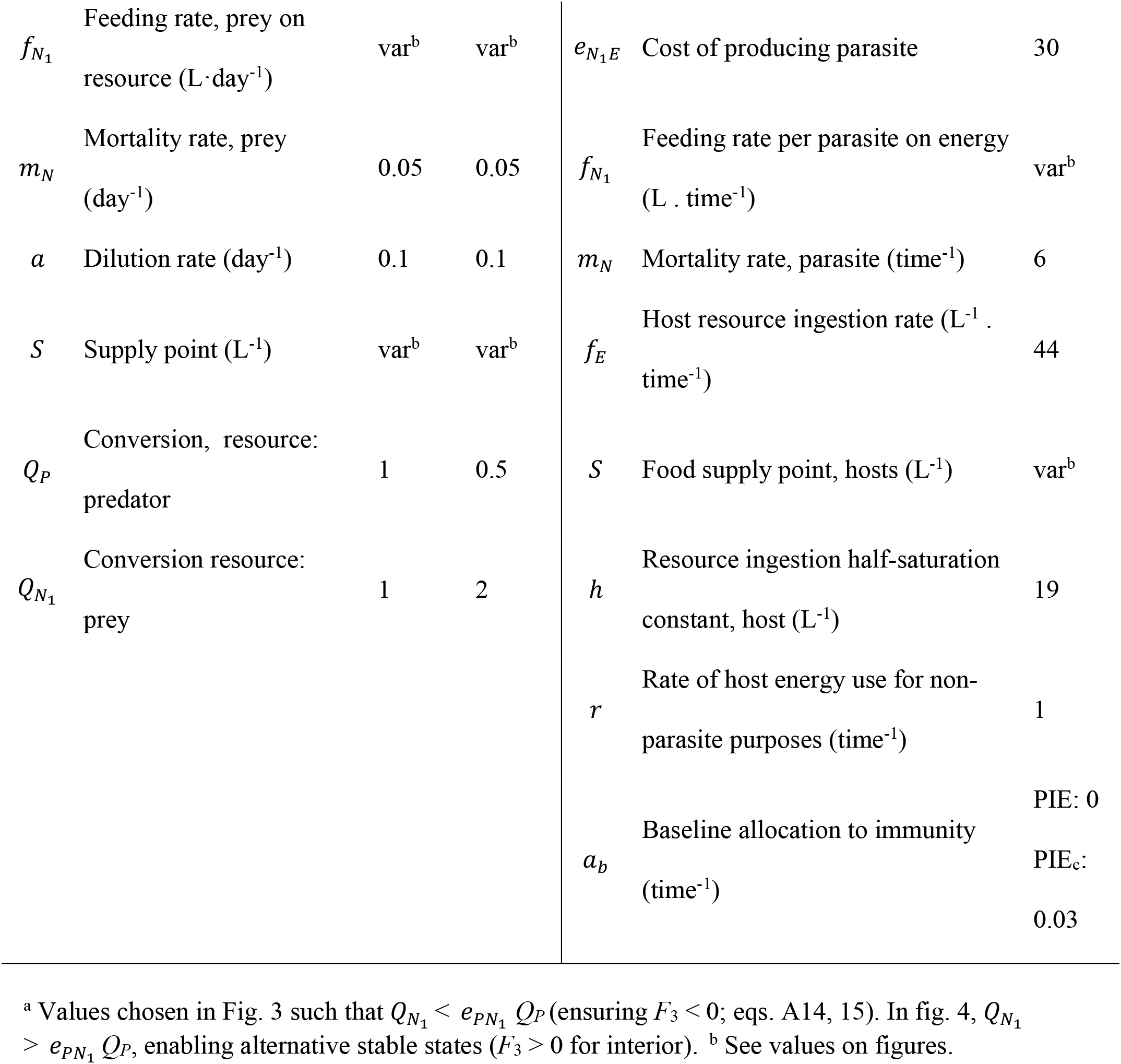
Equations for IGP (Eq. 1) and PIE models (PIE and PIEc constitutive immunity; Eq. 2) with top consumers (predators or immune cells), intermediate consumers (prey or parasite), and basal resource (or host energy). Listed below each model are the variables and parameters, their definition (with units), and corresponding default values used (in Figs. 3,4).

### Intraspecific direct and indirect effects: an approach using feedback (Box 1 and Fig. 2)

To compare stability of these models, we used feedback loops (Box 1). Any *n*-dimensional system can be decomposed into *n* levels of feedback that determine stability of an equilibrium (Box 1, Top Panel). An interior equilibrium can allow stable or unstable coexistence (oscillations) or produce, potentially, alternate stable states (if it is a saddle). As illustrated by and applied to IGP and PIE, the three-dimensional system creates three levels of feedback (Box 1, Fig. 2). For example, level 1 feedback (*F_1_*) is the sum of intraspecific direct effects (Box 1) and is the Jacobian’s trace (Appendix Section 1). At the interior of IGP and PIE, the basal resource / energy contributes to the direct effects at the feasible interior (Fig. 2, Appendix Section 2). Self-limitation comes from the chemostat-supplied resource and its consumption; in PIEc, baseline allocation to immunity adds self-limitation (hence negative level 1 feedback [see below but also Appendix Section 1]). Level 2 (*F_2_*) feedback sums up two-species loops (Box 1). In PIE models (Fig. 2), this sum of pairwise consumer resource come from energy-parasite (*E-N_1_*), energy-immune cells (*E-I*), and parasite-immune cells (*N_1_-I*, with *R*-*N*_1_, *R-P*, and *N*_1_*-P* analogies in IGP, respectively). At the interior, all of these loops contribute to negative feedback. That negative feedback intensifies in PIEc from the product of energy and immune self-limitation. In all 3D models considered here, *F*_2_ is negative at the interior equilibrium (based just on signs alone): level 2 feedback stabilizes.

Level 3 feedback (*F_3_*) sums three-species loops. It is also the negative of the determinant of the Jacobian, - |**J**| (Appendix Sections 1, 2). In PIE models, these loops at the interior are (Fig. 2, *L-R*): (i) “*N_1_ is eaten*”, a negative loop coming from the stabilizing product of the immune-parasite loop and energy self-limitation; (ii) “*starving the enemy*”, a positive loop where a small increase in the parasite reduces energy for the immune cells, hence reduces immune attack (i.e., ↑*N*_1_ →↓*E* →↓*I* → ↑*N*_1_); (iii) “*feeding the enemy*”, a negative loop where a small increase of the parasite 1 stimulates the immune cells which consume energy that then starves the parasite (↑*N*_1_→↑*I* →↓*E*→ ↓*N*_1_). Thus, negative feedback operates when *N*_1_ hurts itself by “feeding” the immune cells. Similar loops arise in IGP. Finally, in only PIEc, (iv) a negative “*N_1_ eats*” loop arises from the stabilizing product of energy consumption by the parasite and immune self-limitation.

Combined, these three levels of feedback determine stability of the interior equilibrium. First, stability requires that each level of feedback is negative (so, *F_1_* < 0, *F_2_* < 0, and *F_3_* < 0). When *F*_3_ < 0, the interior is stable; when *F*_3_ > 0, it is a saddle. Additionally, stability requires that *F_1_ F_2_ + F_3_* > 0, or that upper-level feedbacks (*F_3_*) not be too strong relative to lower-level feedback (*F_1_ F_2_*). All of these conditions have analogies to the Routh-Hurwitz criteria, differing only in sign (Puccia & Levins 1991). Here, we also reframe stability analysis as sums of intraspecific direct effects (*F_1_*, hereafter DE) and of intraspecific indirect effects (*-F_2_* / *F_3_*, the trace of the inverse Jacobian, IE: see Appendix Section 1, Box 1). Written this way, IE is a ratio of the longer loops, where each species interacts with other species in feedback at level 2 (*F_2_*) and level 3 (*F_3_*). It is also the sum of intraspecific indirect effects of each species *N_i_* (lying on tr[-**J**^-1^]; Eq. 5):

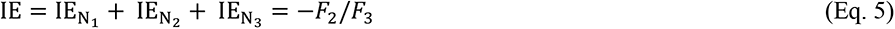

The numerator of the intraspecific indirect effects of each species (IE_Ni_) corresponds to specific combinations of the reciprocal two-species *F*_2_ loops; the denominator for each species is *F*_3_. For example, in general form, the corresponding indirect effect of *N_1_*, IE_N1_, is the sum of (i) the product of self-limitation of *N_2_* and *N_3_* and (ii) pairwise consumer-resource loop of *N_2_* and *N_3_*, all divided by *F*_3_ (green shading over gray shading, Box 1). Similar expressions can be derived for species 2 (IE_N2_, yellow shading) and 3 (IE_N3_, blue shading). With DE and IE in hand, stability then depends on their sign and the strength of their product. Specifically, *stable coexistence* occurs when direct and indirect effects are negative (DE < 0, IE < 0) and jointly strong (DE x IE > 1; bottom of Box 1, light orange region; equivalent to *F*_1_ *F*_2_ > -*F*_3_). *Oscillations* arise when direct and indirect effects are jointly weak (dark orange, DE x IE < 1 for DE < 0, or DE > 0; equivalent to *F*_1_ *F*_2_ < -*F*_3_). Finally, *alternate stable states (aka priority effects)* occur with positive indirect effects (IE > 0, *i.e*., when *F*_3_ > 0 for IGP and PIE, given that *F*_2_ < 0 for both at their interiors; Box 1, gray region). Note: we use the terms alternate stable states and priority effects interchangeably. They both denotes how initial densities, even offset in time (*e.g.,* sequential infection), can determine competitive outcomes (depending on whose domain of attraction they move through).

### Weakening DE x IE generates oscillations in these models (Fig. 3 and Appendix Section 2)

#### Assembly of the interior equilibrium

To compare stability outcomes across IGP and PIE models, we created parallel 2D bifurcation diagrams (Fig 3). These diagrams spread gradients of parameter space of an environmental factor, supply of resource (*S*), and trait space of the focal prey/parasite, feeding rate of *N*_1_ on resource, *f*_*N*_1__ (Fig. 3A-C; interpreted with 1D bifurcation diagrams: Fig. 3G-I). The lines within the 2D bifurcation diagram represent shifts in species composition or in dynamics (stable coexistence, oscillations, or alternative stable states; see Appendix Section 2 for full analysis). In IGP (Fig. 3A), when resource supply and feeding rate of prey are too low to meet the minimal resource requirement of either the prey or predator, only *R* is supported in the system (*R*, yellow). Increases in *S* allows *R* to meet the minimum resource requirement (*R*\*) of the predator (hence *P* invades creating the *R-P* region, green). As feeding rate of prey increases along a fixed resource supply (*S* = 110), the prey becomes a better competitor for resources, enabling births, *f*_*N*_1__*R*\*, to exceed deaths, *f*_PN_1__*P*\* + *m_N_* at the *R-P* boundary. Hence, the prey can invade; both prey and predator push *R*\* lower in the *R-N*_1_-*P* interior (Fig. 3A, Fig. 3G). In this region, *R-N_1_-P* are able to coexist either stably (light orange) or via oscillations (dark orange). On the other hand, as the feeding rate of prey increases at low *S* (*S = 5*), prey invade the resource-only boundary (creating the *R-N_1_* region, blue). The omnivorous predator can then invade this *R*-*N*_1_ boundary when its birth rate, *e_PN_*_1_ *f_PN_*_1_*N*_1_*+ *f*_PR_*R*\*, exceeds its death rate, *m_P_*. Successful invasion creates stable coexistence (orange; Fig. 3A). Thus, *R, R-P, R-N_1_,* and *R-N_1_-P* equilibria (oscillatory and stable coexistence) are possible.

**Figure 3:**
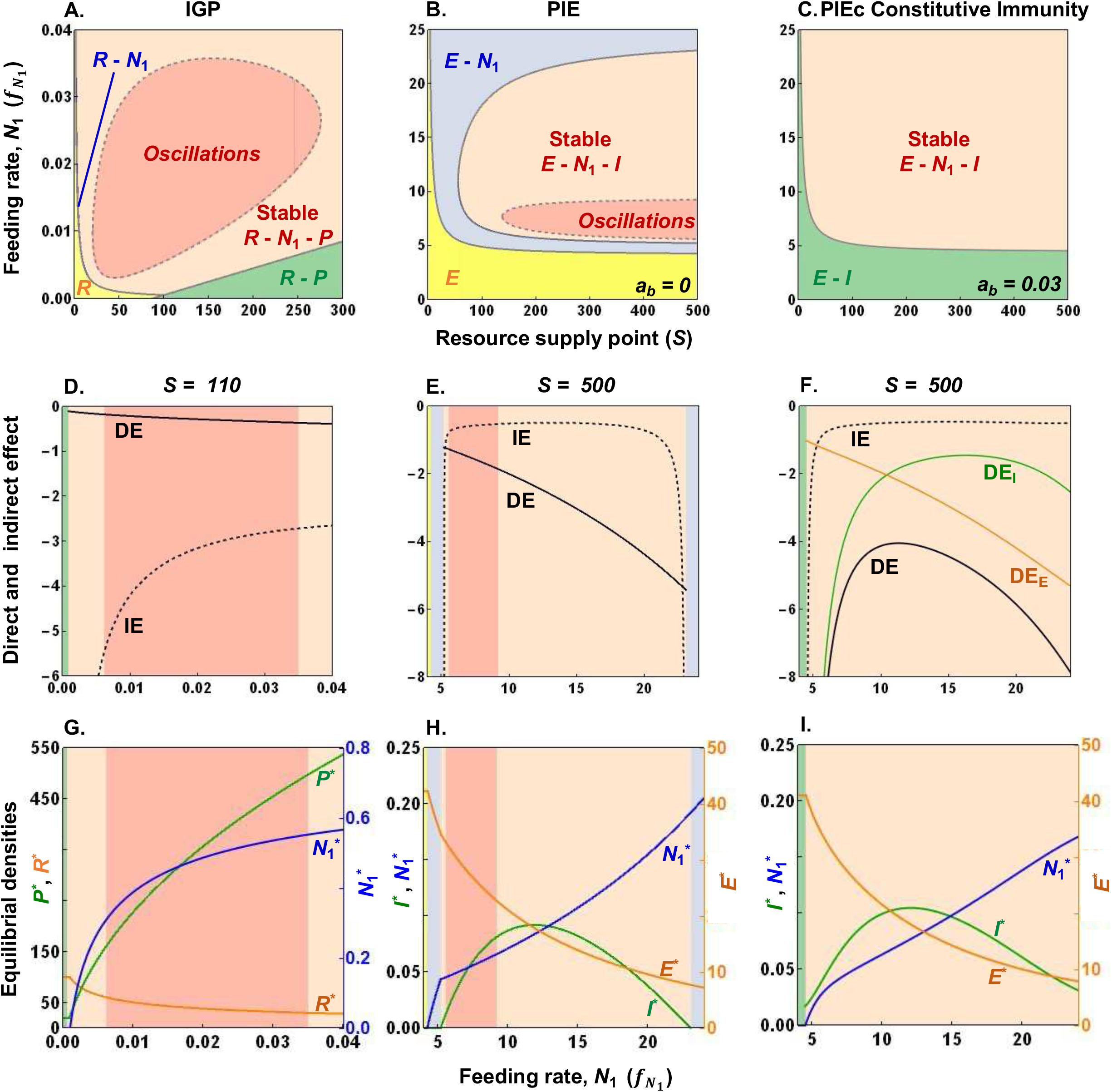
Weakening of the product of intraspecific direct and indirect effects generates oscillations in both IGP and PIE / PIEc models (Fig. 1, Box 1). (A)-(C) *Bifurcation diagrams* over gradients of supply point (*S*) and feeding rate of prey or parasite (*f*_*N*_1__): Oscillations (dark orange region) found in (A) IGP also arise in (B) PIE models of within-host dynamics (see Appendix Section 2). These oscillatory regions are enveloped within a region of stable coexistence (light orange). In PIE models, *a_b_* denotes baseline energy allocated to immune cells: *a_b_* > 0 in (C) PIEc. (D)-(F) *Direct and indirect effects (after* Fig. 2): Oscillations occur with weakening of the product of intraspecific direct effects (DE) and intraspecific indirect effects (IE). Weakening of IE primarily triggers oscillations at low *f*_*N*_1__while strengthening of DE restores stability at higher *f*_*N*_1__in (D) IGP and (E) PIE. In (F) PIEc, self-limitation comes from energy (DE_E_, orange) and immune cells (DE_I_, green). Summed DE eliminates oscillations while lowering parasite burden relative to PIE (see also Fig. A2). (G)-(I) *Equilibrial densities along f_N_*_1_: resource, *R\** or energy, *E\** (orange); prey or parasite, *N_1_\**, (blue); and predator, *P\** or immune cells, *I\** (green) in (G) IGP, (H) PIE, and (I) PIEc models. Shifts in these densities weakens or strengthens DE and IE (see text and Appendix Section 2 for details; Table 1 for default parameters).

The within-host PIE model contains some but not all analogous states. The key differences stem from how hosts produce immune cells. In PIE (Fig. 3B), immune cells only generate with parasites. Hence, a host energy-immune (*E-I*) state, analogous to the *R-P* one, is not possible. Similar to IGP, though, low feeding rate can prevent parasite invasion (creating the *E*-only state [yellow]). With increasing feeding rates (illustrated at *S* = 500), parasites (*N*_1_) can eventually invade creating *E-N_1_* space (light blue; Fig. 3H). Although parasites depress *E*\*, sufficiently high parasite density enables production of immune cells, when the immune system’s minimal *E N*_1_ requirement is met (Fig. 3H). Hence, at medium to high feeding rates of parasites, *E-N_1_-I* coexist either stably (light orange) or via oscillations (dark orange). However, at sufficiently high *f*_*N*_1__, parasites compete too strongly: they successfully starve out the immune cells, re-establishing a *E-N_1_* space (blue). Thus, PIE exhibits only three equilibria: *E, E-N_1_,* and *E-N_1_-I* (oscillations and stable coexistence; Fig. 3B; see Appendix Section 3B for details).

In PIEc, the host always allocates energy to the immune system (*a_b_* > 0). This biology eliminates the *E*-alone and *E-N_1_* region. Instead, immune cells prevent infection (*E-I* space; low *f*_*N*_1__; green) or hosts become infected (*E-N*_1_-*I* space; higher *f*_*N*_1__; orange). Parasites successfully infect when their births, (*f*_*N*_1__/*e*_*N*_1__/_*E*_) *E*\*, exceed their deaths at immune-enhanced, *m_I_* + *f*_*1N*_1__*I*\*. Like IGP, at high feeding rates of parasites, the *E-N_1_-I* remains stable (light orange) or oscillates (only at low *a_b_* [Appendix Section 2]). Thus, in PIEc, immune biology eliminates even more states; only *E-I* and *E-I-N_1_* (oscillations and stable coexistence) remain possible (Fig. 3C).

### Stability outcomes

Despite these differences, oscillations arise for similar reasons in the food web and within-host models. As noted in Box 1, oscillations require weakening of the joint product of intraspecific direct effects (DE) and indirect effects (IE). In all three cases, stable coexistence (DE x IE > 1) envelopes oscillatory regions (DE x IE < 1; Fig. 3A-B; Appendix Fig. A2). In IGP, weakening of IE at low feeding rate of the prey, *f*_*N*_1__, triggered oscillations despite strengthening of DE (Fig. 3D). At high *f*_*N*_1__, DE strengthens to regain stability. This general pattern also arises with increasing feeding rate of parasites in the PIE and PIEc models (Fig. 3E-F). However, DE differed slightly among models (Fig. 3D-F, black, solid lines), where DE is:

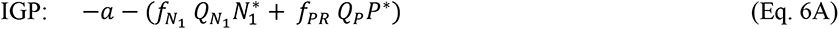

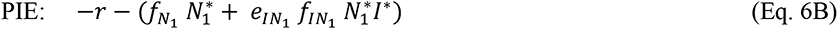

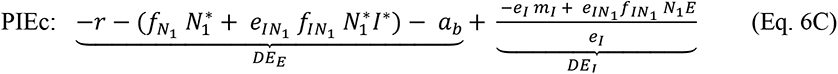

Here, either the sum (IGP) or sum and product (PIE) of victim and enemy densities strengthens DE (Eq. 6). Strong enough DE then tips the coexistence equilibrium from oscillations to stability. In IGP the additive increase in predator and prey densities strengthens DE of resources (Eq. 6A). Thus, strengthening of DE restores stability at higher *f*_*N*_1__ (Fig. 3D, black, solid line). In the PIE models, however, increase in the (weighted) sum of parasites plus the product of parasite and immune cells strengthens DE of energy at higher *f*_*N*_1__ (Eq. 6B-C; Fig. 3E-F, black, solid line).

Constitutive immunity enhanced stability in PIEc models by creating immune self-limitation. This self-limitation generates additional loops featured in both intraspecific direct effects (DE) and indirect (IE; Fig. 2). In PIEc, DE now has two components: DE of energy (DE_E_) and DE of immune cells (DE_I_; Eq. 6C). The updated DE_E_ adds fixed energy allocation towards baseline immunity (–*a_b_*; Fig. 3F, orange, solid line [DE_E_]). DE_I_ is always negative, too (see Appendix Section 2C). This added immune self-limitation (Fig. 3F, green, solid line [negative DE_I_]) further strengthens DE (as shown in Fig. 3E: black, solid line), potentially enough to help to eliminate even the possibility of oscillations in PIEc (eliminated in Fig. 3C; see Appendix Section 2C, Fig. A2). Constitutive immunity also elevates density of immune cells (*I*\*), lowering parasite burden (*N*_1_*) while maintaining slightly higher *E*\* available to hosts for metabolic work (at *rE*\*; compare Figs. 3H vs. 3I). Hence, constitutive immunity both stabilized dynamics via self-limitation in PIEc (Fig. 2) and reduced parasite burden.

At low feeding rate in both IGP and PIE models, weakening of IE of the prey or parasite triggers oscillations. IE is a ratio of shorter, binary loops (*F_2_*) and longer, three-species loops (*F_3_*) (Fig. 2). In all three models, *F_3_* became more negative with feeding rate of the prey/parasite, *f*_*N*_1__, than did *F_2_* (as parameterized). That helps explain why the ratio *–F_2_*/*F_3_* (IE) became less negative with increasing *f*_*N*_1__– and hence why weakening IE triggered oscillations (Figs. 2, 3D-F). However, the precise details varied among the models (so it becomes unfruitful to generalize here).

### Positive feedback leads to alternate stable states in IGP alone (Fig. 4 and Appendix Section 2)

Despite the parallels described above, IGP model can also produce alterative stable states not found in the with-host analogues. In fact, using different parameter values (Fig. 4A; see Appendix Section 2A), a 2D bifurcation diagram shows two forms of alternate stable states in IGP (Fig. 4A; Alt-SS1 [dark grey] and Alt SS-2 [light grey]; see also Verdy & Amarasekare 2010). In these regions, initial densities of species determine competitive outcomes (Fig. 4C, D). The simpler form of alternative stable states (Alt SS – 1; Fig. 4C) yields just the prey (*R-N_1_*) or just the predator (*R-P*) with the resource. If it held in PIE, analogous outcomes would lead to exclusion of immune cells by parasites (yielding stable infection in the *E-N*_1_ state) or vice versa (yielding immune clearance, a stable *E-I*), depending on initial parasite dose. However, in PIE, the host cannot generate immune cells without parasites. Consequently, since the *E-I* state is both biologically and mathematically not feasible, the analogy breaks (Appendix Section 2A). In the other type of alternative stable states in IGP (Alt SS – 2; Fig. 4D), we find either coexistence (*R-N_1_-P*) or a predator-only (*R-P*) equilibrium, depending on initial conditions (due to two feasible interior equilibria, one of which is a saddle). Such outcomes would resemble a host that became infected (*E-N_1_-I*) or cleared infection (*E-I*), depending on parasite dose. We could not fine such an outcome in PIEc (see Appendix Section 2C). Hence, despite the qualitatively similar loop structure, IGP offered two scenarios for alternative stable states not present in within-host analogues. Alternative infection states would require some other mechanism to produce positive feedback.

**Figure 4:**
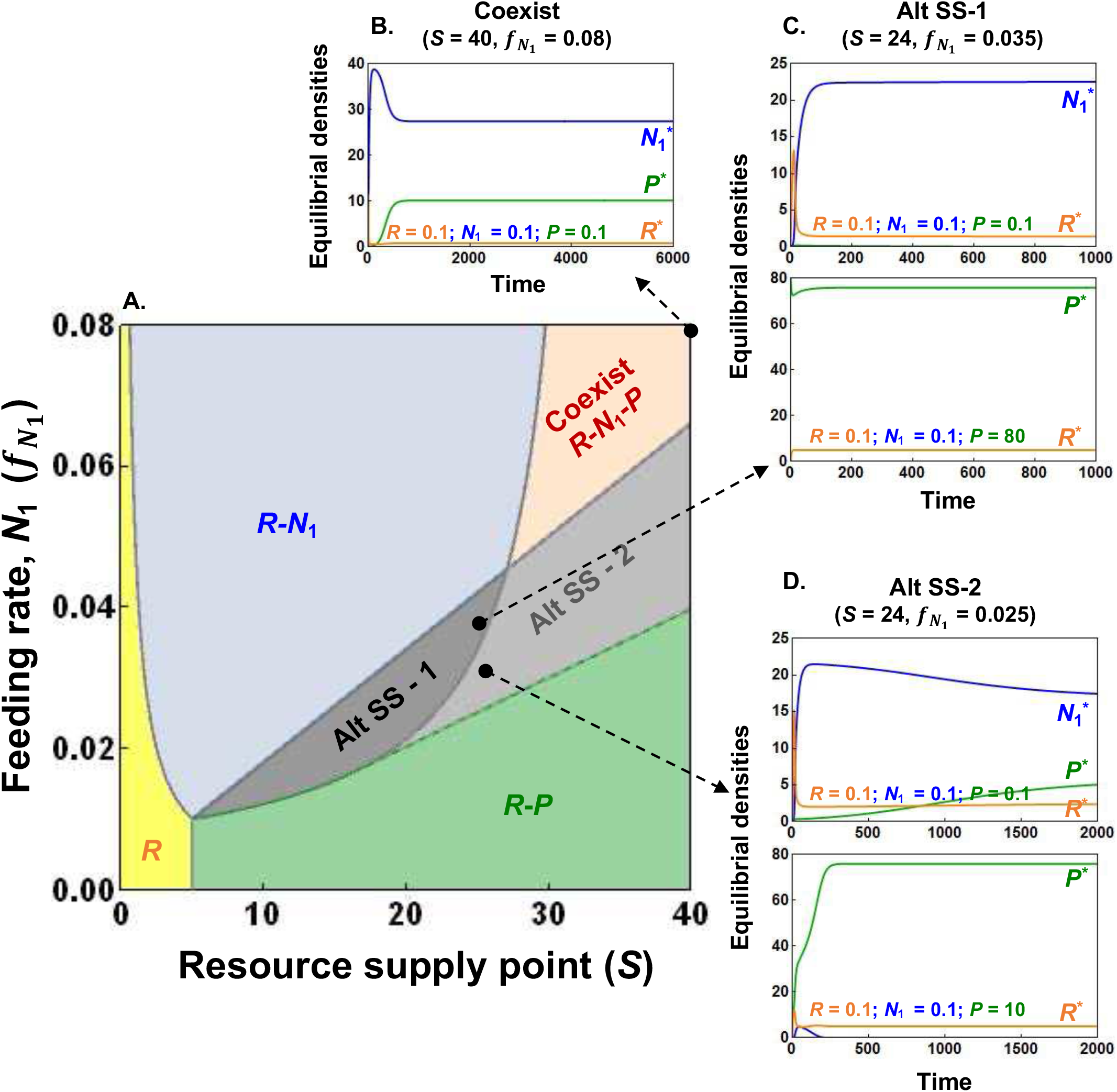
Positive intraspecific indirect feedback leads to two forms of alternate stable states (Alt SS-1 and -2) in the IGP model of prey (N_1_), predators (P), and resources (R) not found in PIE models. (A) A bifurcation diagram over gradients of resource supply point (*S*) and prey feeding rate (*f*_*N*_1__) and species dynamics at the interior equilibrium (see Fig. A2 for more details). (B-D) Sample dynamics. (A,B) Stable coexistence (*R – N_1_ – P,* light orange) found in IGP also arises in PIE predicting successful infection (Fig 3; *E – N_1_ – I*). (B-D) IGP predicts two forms of alternate stable states, where initial densities (inset) determine competitive outcomes between *N*_1_ and *P*. (A, C): In more typical alternative stable states (Alt SS-1; dark grey), either prey (*R*-*N*_1_) or predator (*R-P*) wins. Analogous within-host states would lead to infection (*E-N_1_*) or not (*E-I*). (A, F) Another example (Alt SS – 2; light grey) predicts either coexistence (*R*-*N*_1_-*P*) or an only-predator (*R-P*) equilibrium (analogous to infection [*E-N_1_-I*] or clearance [*E-I*]). Neither of these types of alternative states emerge in PIE models (see text, Appendix Section 2).

### Two prey – predator - resource (KP) | Two parasite - immune cells - host energy (2PIE / 2PIEc)

#### Model Summary (Fig. 1, Table 2, Appendix Section 3)

Both keystone predation (KP) and within-host (2PIE) models feature two competitors engaged indirectly via consumer-resource like interactions. In both, consumers directly benefit (positive, black arrow) with direct costs to resources (negative, red circle). In KP, predators (*P*) can consume two prey (*N_i_*), leading to positive interspecific direct effects for the predator and negative effects on prey (Fig. 1D). The two prey themselves compete for a shared resource (*R*), creating similar +/- interactions. Additionally, the resource experiences self-limitation at the interior (four species) equilibrium (Fig. 1D). Unlike in IGP, the predator does not consume the resource (i.e., it is not omnivorous). Hence, competitors engage only in apparent and exploitative competition (with *P* and *R*, respectively). Similarly, 2PIE and 2PIEc extend the within-host PIE models. Now, two parasites (*N_i_*) compete for host energy but are attacked by immune cells (Fig. 1E-F). During these attacks, immune cells use host energy to proliferate (as described in the PIE models). In 2PIEc, hosts also directly allocate energy to production of immune cells (*a*_b_ > 0), forging new connections and self-limitation of immune cells (Fig. 1F, Appendix Section 2). Further, *E-I* links introduce additional components of loops not found in KP (e.g., in *F_3_*; Fig 5). Therefore, it might seem that KP provides too simple of a comparison to 2PIE/2PIEc. In fact, 2PIE models best resemble a combination of KP and IGP. Yet, the feedback loops that determine coexistence versus alternate stable states (*F_4_*) of the prey or parasite are qualitatively similar (Fig. 5). The resemblance arises because both the prey and parasite are similarly *effected by* and have *affects on* their resources and their enemy. However, stronger negative feedback at level 3 in 2PIEc will shrink opportunities for coexistence while lowering parasite burden (see below).

**Figure 5:**
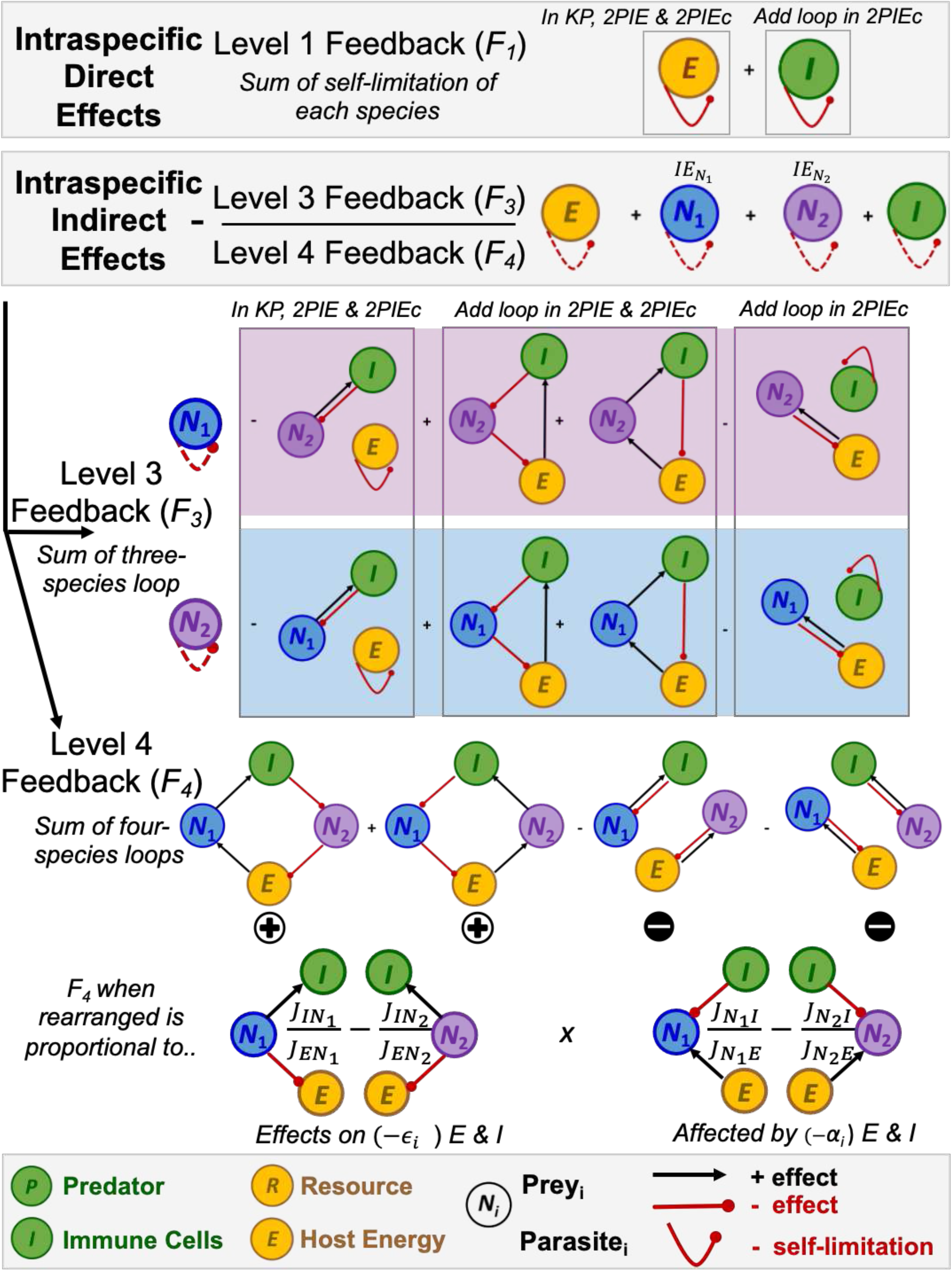
Intraspecific direct and indirect effects in four-dimension models: feedback in KP, PIE, and 2PIEc (Fig 1). Evaluated at a feasible interior equilibrium, feedbacks for KP and 2PIE are nested within 2PIEc. In 2PIE and 2PIEc, additional loops arise from interactions between immune cells and host energy (“add loop in 2PIE & 2PIEc”), and immune self-limitation loop (“add loop in 2PIEc”). Level 1 feedback (*F_1_*) sums intraspecific direct effects (solid curves) from energy and immune self-limitation (-; only in 2PIEc). The intraspecific indirect effects (dashed curves) involve a ratio of longer loops, where each species, *N_1_* (IE_N1_; blue shading) and *N_2_* (IE_N2_; purple shading), interacts with others in feedback at level 3 (*F*_3_) and 4 (*F*_4_). *F_3_* sums three species loops in PIE models: (i) “*N_1_ is eaten*”, the *I-N_1_* loop times *E* self-limitation (-; all models), (ii) “*starving the enemy*” (+; from PIE models), (iii) “*feeding the enemy*” (-; from PIE models), and (iv) “*N_1_ eats*”, the *E-N_1_* loop times *I* self-limitation (-; only from PIEc). *F_4_* sums, from *L-R*, two destabilizing (+) loops and stabilizing (-) ones. *N_i_* benefits as it (i) “*feeds*” or (ii) “*starves*” the enemy, but is restrained by (iii) and (iv) “*N_i_ is eaten, N_j_ eats*” loops (the product of *I-N_i_* and *N_j_-E* loops). With some rearrangement, *F_4_* becomes proportional to differences in ratios of how each competitor has *effects on* (*ε*_1_ – *ε*_2_) and is *affected by* (*α*_1_ – *α*_2_) immune cells and host energy (see text). Finally, the sign of *F_4_* determines that of IE (see text). The predator (*P*, green), prey 1 (*N_1_*, blue), prey 2 (*N_2_*, purple), and resources (*R*, yellow) in KP are analogous to immune cells (*I*), parasites (*N_1_* and *N_2_*), and host energy (*E*) in 2PIE models, respectively.

#### Intraspecific direct and indirect effects in the 4D KP and 2PIE model (Fig. 5)

Similar to 3D systems, the sign of intraspecific indirect effects determines coexistence vs. priority effects in 4D systems. But first, in both KP and 2PIE, only self-limitation of the resource or host energy, respectively, contribute to summed intraspecific direct effects (DE; level 1 feedback, *F_1_*; Fig. 5). 2PIEc has additional contributions from the immune self-limitation. Then, summed intraspecific indirect effects (IE) involve a ratio of feedback loops with three species (*F_3_*) and four (*F_4_*; IE = -*F_3_*/*F_4_*; Fig 5; Appendix Section 3D). In KP and 2PIE models, *F_3_* sums the negative consumer-resource loop between each victim and its enemy times self-limitation of the resource (analogous to loop [i] in PIE [Fig. 2]). Then, energy-immune interactions add two sets of additional loops, analogous to positive “*starving the enemy*” and negative “*feeding the enemy*” loops in PIE, but for each parasite (loops [ii] and [iii], respectively). Finally, immune self-limitation in 2PIEc adds two more negative feedback loops (involving *N_i_ – E* for each parasite; loops [iv]). At the interior, we find negative *F_3_* for the interiors evaluated here (algebraically or numerically). Additionally, the sum of loops involving parasite *N*_2_ provide the numerator of the intraspecific indirect effect of *N*_1_ (IE_N1_; blue shading), while those involving *N*_1_ provide that of *N*_2_ (IE_N2_ : purple shading; Fig. 5). Unlike in IGP/ PIE, the intraspecific indirect effect of the enemy and resource now becomes zero. Hence, only the sum of IE_N_i__ determines stability: coexistence occurs when each competitor *N_i_* indirectly decreases its own growth rate.

Importantly, level 4 feedback loops are qualitatively similar across the KP, 2PIE, and 2PIEc models. These loops (from *L-R*) are: (i) a positive *“feeding the enemy”* loop, where a small increase in one parasite stimulates the immune cells which attack the competing parasite, freeing up energy for the first parasite. In a second positive loop, (ii) *“starving the enemy”,* a small increase in a parasite reduces resources, hence density of the competing parasite, thereby lowering immune activation, ultimately reducing mortality on the first parasite. Those two positive (destabilizing) loops then push against two negative (stabilizing) loops, (iii and iv) “*N_i_ is eaten, N_j_ eats*”. These loops therefore add the stabilizing consumer-resource interactions within which each prey/parasite is enmeshed. For instance, a small increase in parasite *i* creates a negative loop with the immune system while parasite *j* is braked by its loop with the resource. Then, those roles reverse, i.e., parasite *j* is slowed by the immune system and parasite *i* is by the resource. Summed, those two loops (iii and iv) jointly determine the amount of negative feedback in the system at level 4. Combined then, loops (i)-(iv) determine the sign of *F_4_*, the sign of the summed intraspecific indirect effects (*-F_3_*/*F_4_*; since *F_3_ <* 0: see above), and stability of a feasible interior equilibrium.

In these 4D systems, stability of the interior can also be understood via symmetries or asymmetries in two quantities. These quantities emerge upon rearranging the loops comprising the level 4 feedback (*F_4_*). They reflect ratios of how each prey or parasite has effects on (-*ε_i_*) and is affected by (-*α_i_*) their enemies and resources (Fig 5; Eq. 7):

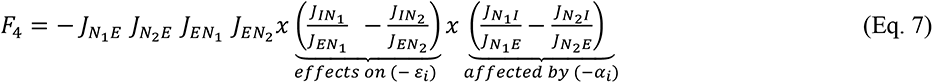

where *J_ij_* are interspecific direct effects (Jacobians) of species *j* on species *i*, 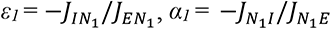, *etc.* Stability hinges on (a)symmetry of these ratios. As described elsewhere (Appendix Sections 3A, B), the difference in *affected by* ratios ensures a tradeoff in traits influencing resource and apparent competition that permits coexistence. If *N*_1_ is the superior resource competitor without the enemy, then *α*_1_ > *α*_2_ ensures *N*_2_ is sufficiently resistant to attack. The difference in *effects on* ratios then determines arrangements of competitive hierarchies with enemies (involving exploitative [resource, energy] and apparent [predator, immune] competition). Coexistence occurs with symmetry in these ratios (i.e., if *α*_1_ > *α*_2_ if *ε*_1_ > *ε*_2_), guaranteeing that net negative feedbacks dominate (*F_4_* < 0). In contrast, asymmetry in these ratios, (e.g., *α*_1_ > *α*_2_ still but now *ε*_1_ < *ε*_2_) leads to net positive feedback (*F_4_* > 0), triggering alternate stable states.

#### Coexistence (coinfection) and alternate stable states (priority effects) in KP & 2PIE (Fig. 6 and Appendix Section 3)

Despite differences in biology, KP and 2PIE models share similar four species feedback loops, hence qualitatively similar outcomes for stability of the interior equilibrium. That similarity becomes readily apparent in 2D bifurcation diagrams for each model in parameter space of supply point (*S*) – *N*_1_’s feeding rate on resources (*f*_*N*_1__). In KP (Fig. 6A), at low *S-f_N_*_1_, only the resource is supported in the system (*R*, yellow). Increased feeding rate at low resource supply enables prey *N_1_* to meet its minimum resource requirement (*R-N_1_*; light blue). As *S* increases, the minimum prey requirement of the predator is met (its *N_1_*\*) allowing predators to invade (*R-N_1_-P;* dark blue). Then, the more resistant *N*_2_ can invade when the *R*-*N*_1_-*P* food chain provides enough resources, given predator density (mortality). This invasion enables four species coexistence (*R*-*N*_1_-*N*_2_*-P;* light orange). Similar assembly rules apply for invasion of *N_2_* when feeding rate of *N*_1_ stays low enough to grant *N*_2_ competitive superiority (*R*_2_* < *R*_1_*). With increasing *S*, first *N*_2_ invades (*R-N_2_* space; light purple), then the predator does (*R-N_2_-P*: dark purple). A less typical assembly arises at lower *S* (*S = 1*) but with increasing *f*_*N*_1__. Here, *N*_1_ and *P* simultaneously invade creating the jump from 2 (*R-N_2_*) to 4 (*R*-*N*_1_-*N*_2_*-P*) dimensional stability (see Appendix Section 3C for details). Overall, similar transitions appear in 2PIE models. Like KP, the possible states in 2PIE models are energy alone (*E*), just one parasite (*E-N_i_*), addition of immune cells (*E-N_i_-I*), or an interior (*E-N_1_-N_2_-*I; Fig. 6B). However, since hosts always allocate energy to immune cells, they are always present in 2PIEc. Hence, 2PIEc only has *E-I, E-I-N_i_*, and *E-N_1_-N_2_-I* states (but not *E* or *E-N_i_*; Fig. 6C).

**Figure 6:**
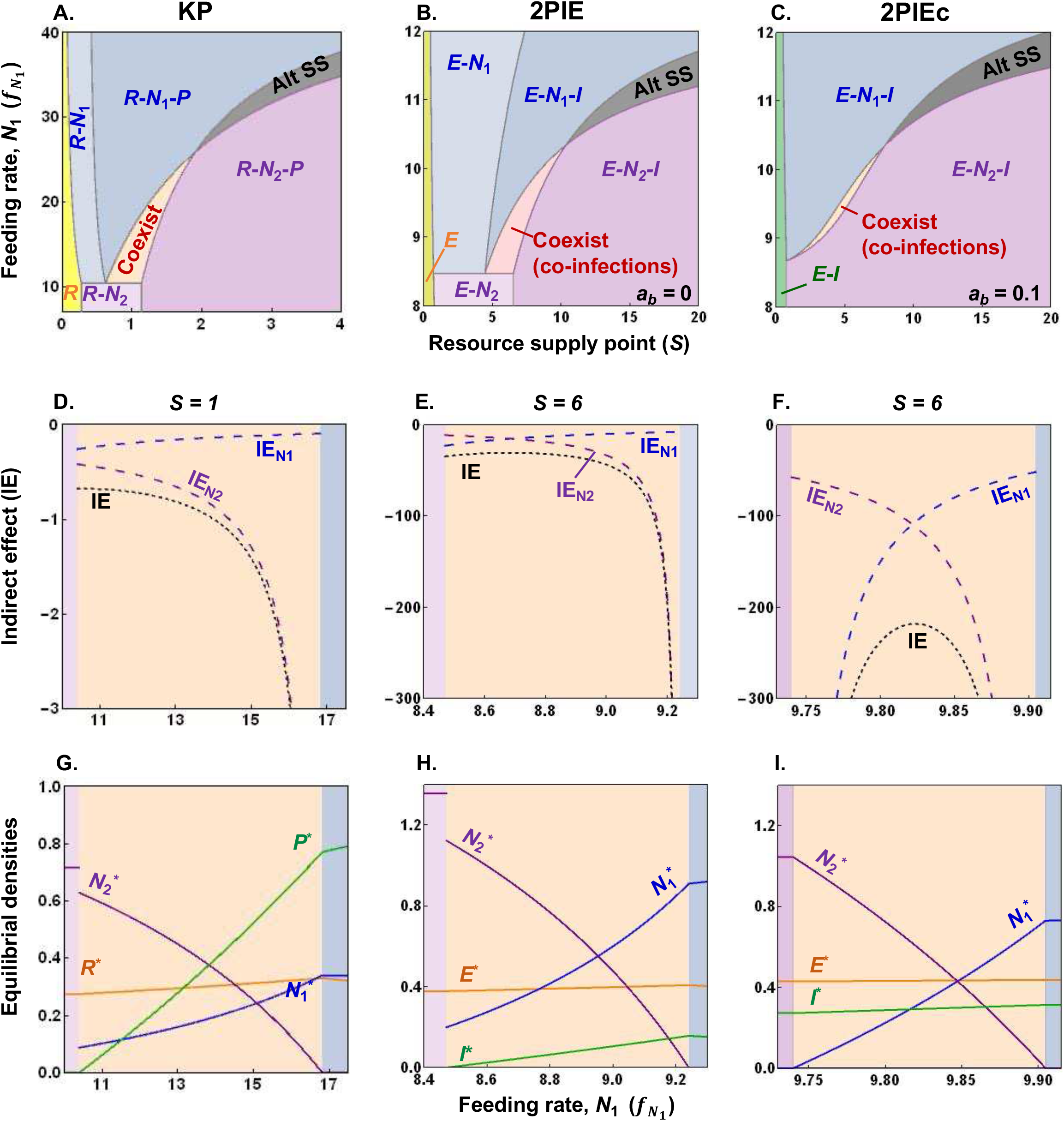
KP anticipates coexistence and alternate stable states in 2PIE models. (A)-(C). *Bifurcation diagrams* over gradients of resource supply point (*S*) and feeding rate of prey or parasite (*f*_*N*_1__): alternative stable states or priority effects (grey region) and stable coexistence (orange) found in (A) KP also arise in (B) 2PIE and (C) 2PIE constitutive immunity (2PIEc) models of within-host dynamics. In 2PIE, *a_b_* denotes baseline energy allocated to immune cells. Stable coexistence (coinfection) happens at low *f*_*N*_1__and *S* in all models. (D)-(F) *Indirect effects through coexistence/coinfection regions*: Strong, negative indirect effects (IE; black, short dash) from *N*_1_ (IE_N_1__; blue, long dash) and *N*_2_ (IE_N_2__; purple, long dash) shift systems from coexistence to exclusion of *N*_1_ or *N*_2_ at lower or higher *f*_*N*_1__ respectively in (D) KP, (E) 2PIE, and (F) 2PIEc. (F) Immune self-limitation in 2PIEc enhances negative indirect effects and squeezes parameter space enabling coinfection. (G)-(I) *Equilibrial densities along f_N_*_1_: Resource, *R* or energy, *E* (orange); prey or parasite 1, *N_1_*, (blue); prey or parasite 2, *N_2_*, (purple); and predator, *P* or immune cells, *I* (green) in (G) KP, (H) 2PIE, and (I) 2PIEc models. (See text and Appendix Section 3 for details, Table 2 for default parameters).

**Table 2.**
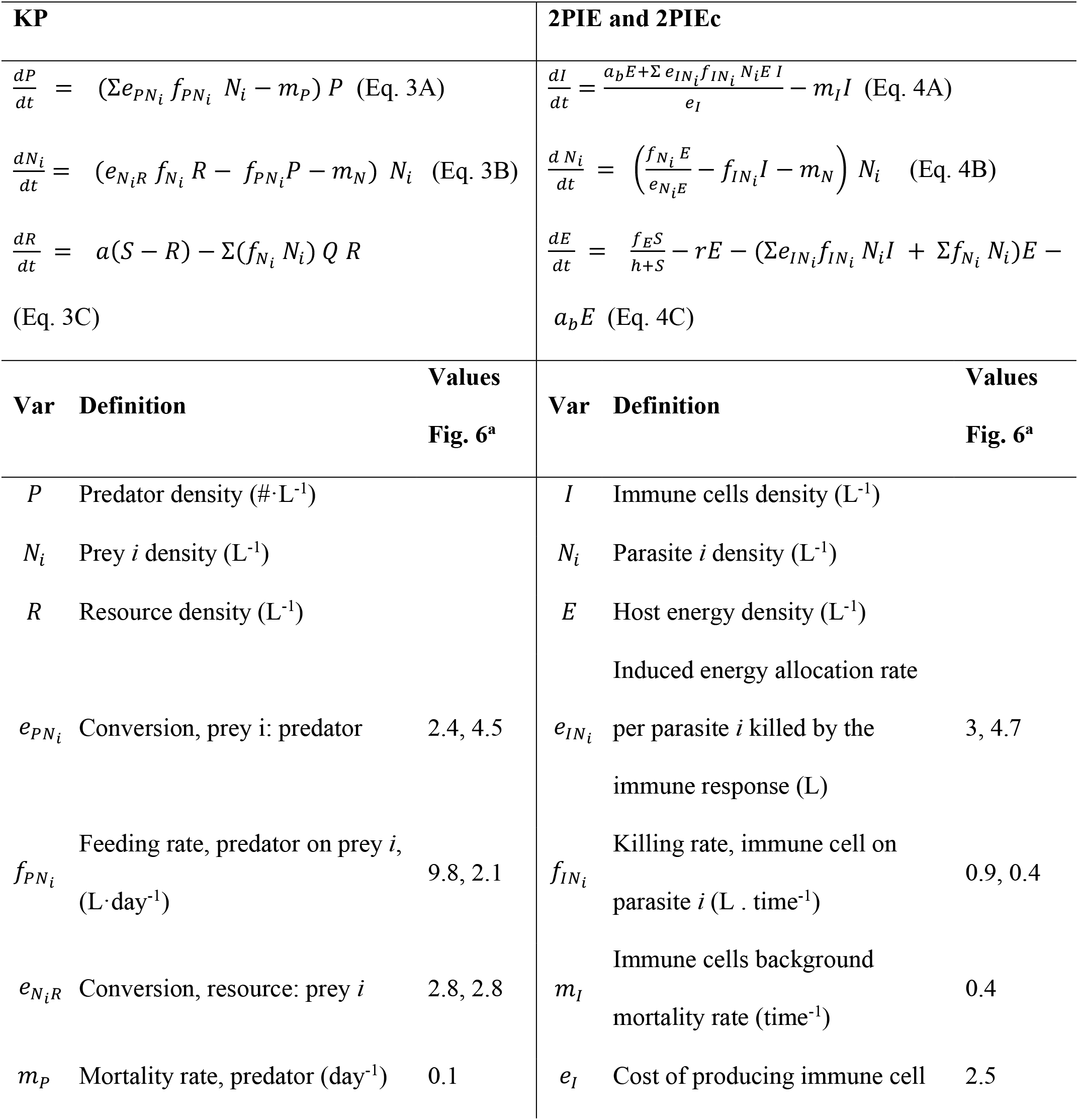
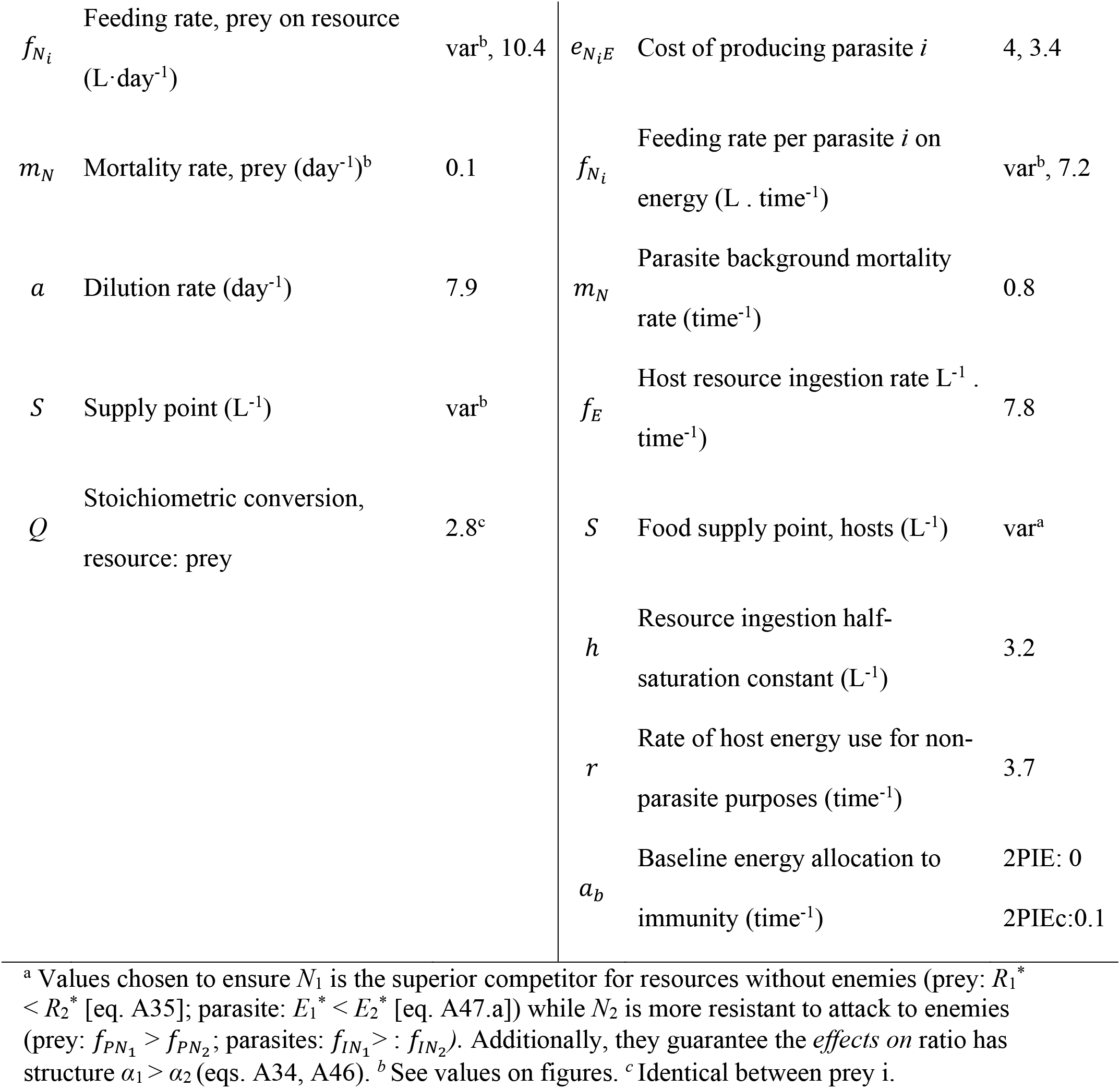
Equations for KP (Eq. 3) and 2PIE models (2PIE and 2PIEc constitutive immunity; Eq. 4) with top consumers (predators or immune cells), intermediate consumers (prey or parasites), and basal resource (or host energy). Listed below each model are the variables and parameters, their definition (with units), and corresponding default values used (in Figs. 6).

In all three models, low or high combinations of supply point (*S*) and feeding rate (*f*_*N*_1__), create (a)symmetry in *effects on* and *affected by* ratios that, in turn, generate coexistence or alternate stable states for a feasible interior. We assume *N*_1_ competes superiorly for energy without immune cells but is sufficiently more vulnerable to immune attack (enabling *α*_1_ > *α*_2_). In the coexistence region (i.e., at lower *f*_*N*_1__*-S*), *N_1_* is the superior apparent competitor while more resistant *N*_2_ is the superior resource competitor (see Appendix Section 3A,B for details and proof). Such a switch in competitive hierarchy means *N*_1_ exerts stronger effects on immune cells relative to energy (*ε*_1_ > *ε*_2_; Eq. 7). It also produces a symmetry in ratios that generates net negative feedback (*F_4_* < 0) and negative intraspecific indirect effects of the competitors (IE*_i_* < 0) enabling coexistence (coinfection; 6D-F). However, high *f*_*N*_1__-*S* produces an asymmetry: *N_2_* now is the superior apparent competitor, and it has stronger effects on immune cells relative to energy (*ε*_1_ < *ε*_2_) while *N*_1_ is the superior resource competitor. This asymmetry in ratios generates net positive feedback (*F_4_* > 0) triggering alternate stable states (or priority effects; Appendix Section 3A,B). Hence, this interior saddle separates dominance by one or the other parasite (or prey). Furthermore, intraspecific indirect effects are both positive (IE_Ni_ >0): each competitor now benefits itself through the feedback loops. With even higher *f*_*N*_1__, *N*_2_ becomes excluded in all three models (Fig. 6G-I). Notice also how, at the border of that exclusion from the coexistence region, IE_N2_ becomes extremely strong (infinitely negative, since *F_4_* = 0; Fig. 6D-F). In other words, the excluded species suffers extremely strong negative intraspecific indirect effects (i.e., IE_N_2__ → -∞).

The 2PIEc model modifies the patterns in 2PIE in qualitative and quantitative ways. Qualitatively, the range of outcomes in *S*-*f*_*N*_1__space simplifies (as in PIEc: Fig. 4C). Since baseline allocation guarantees positive density of immune cells, there are no *E* or *E-N*_i_ regions like in 2PIE (Fig. 6B vs. 6C). Quantitatively, in the co-infection region, constitutive immunity strengthens intraspecific indirect effects (because of the added negative feedback in the *F_3_* loops, the numerator of IE of each species [Fig. 5]). Stronger IE, in turn, narrows the parameter space permitting coexistence. Constitutive immunity also elevates density of immune cells (*I*\*), lowering parasite burden (*N*_1_* + *N*_2_*) while maintaining slightly higher *E*\* (allocated at *rE*\*) for other metabolic work (compare Figs. 6H vs. 6I). Hence, constitutive immunity adds to *F_3_* loop structure of 2PIEc (Fig. 5). That addition squeezes coinfection space while reducing parasite load relative to 2PIE.

## DISCUSSION

What within-host factors determine parasite dynamics? To answer this question, we compare two classic food web modules, intraguild predation (IGP) and keystone predation (KP), to their within-host analogues (PIE and 2PIE). On the one hand, the comparison seems apt. Both prey and parasite consume a shared resource while facing mortality from an enemy (predator or immune cell). On the other hand, predator and immune cells are produced rather differently. For instance, predators can eat only prey or the resource or both (in IGP). In contrast, induced proliferation of immune cells requires both energy and parasites simultaneously. Furthermore, hosts can allocate energy to immunity constitutively (a pipeline uncommon to food webs). Given these similarities and differences, we analyze stability of equilibria produced by each model using feedback loops. In particular, we interpret loops as direct and indirect intraspecific effects, and we show how their sign and strength govern stability outcomes (Box 1). With these tools, we make three points. First, both IGP and PIE systems predict stable coexistence / infection or oscillations due to similar shifts in direct and indirect effects (Fig. 2-3). Second, that enemy-generation difference eliminates alternative stable states (priority effects) seen in IGP – the PIE models cannot produce them without inclusion of some other mechanisms (Fig. 4). Third, despite that KP is structurally simpler than 2PIE models, competing prey and parasites coexist (co-infect) or show priority effects for essentially the same reasons (Fig. 5-6). The outcomes hinge on the sum of similar four-dimensional loops that generate positive and negative feedback. We show how those loops, in turn, translate into parallel symmetries (stabilizing) or asymmetries in *effects on* and *affected by* ratios of the two competitors. Overall, food web models offer powerful if imperfect analogies to feedbacks underlying the dynamical repertoire of parasites within hosts.

Feedback loops enabled biologically meaningful comparisons of the stabilizing and destabilizing forces in the models analysed here. Traditionally, one evaluates (local) stability using Routh-Hurwitz criteria (Routh 1877; Hurwitz 1895). By rearranging these criteria, we unpack the biology of the feedback loops underlying stability (Box 1; Puccia & Levins 1991; Novak *et al* 2016). These loops can involve shorter, direct effects (DE), where increases in intraspecific density leads to self-limitation (negative feedback) or facilitation (positive). Longer loops involve chains of connected interactions – here, between two (e.g., binary consumer-resource), three, or four species (Puccia & Levins 1991). We show that certain ratios of these loops at different feedback levels, in turn, correspond to intraspecific indirect effects (IE – falling along the trace of the inverse Jacobian matrix evaluated at the equilibrium; Box 1). Furthermore, we illustrate how negative IE leads to coexistence (stable or oscillatory [in 3D systems]), and positive IE can produce alternative stable states. In 3D systems, that transition from stability to oscillations involves weakening of DE x IE. Such weakening occurs when feedback at longer loops becomes too strong relative to those at shorter loops. With that framework in mind, the loops then facilitated comparison between structurally similar but biologically distinct systems (Novak *et al* 2016), *e.g.,* food web and within-host modules. Indeed, loops can reveal the biology obscured or overlooked in stability analysis.

### Single infection dynamics

This loop-based approach to stability analysis revealed that single prey and parasite can persist – stably or via oscillations – with their resource and enemy for similar reasons. Generally speaking, like in IGP, coexistence in PIE requires summed negative intraspecific indirect effects (IE). Like in IGP, weakening of IE (-*F*_2_/*F*_3_) triggers oscillations in PIE (due to weakening of *F*_3_ relative to *F*_2_; Fig. 3D-E). Yet, either the sum (IGP) or sum and product (PIE) of victim and enemy densities strengthens intraspecific direct effects (DE). Strong enough DE then tips the coexistence equilibrium from oscillations to stability. Those oscillations matter because they create boom-bust cycles of parasite and immune response that can harm hosts (Otterstatter & Thomson 2006; Childs & Buckee 2015). For instance, these oscillations produce rapid ramp up or down of immune function that can stress host physiology (exacting e.g., oxidative damage, cellular senescence), making them sicker (Costantini & Mølle 2009). However, in hosts with high constitutive immunity (PIEc), the strengthening of DE by immune-self limitation can prevent oscillations, potentially reducing immune damage to host. Additionally, constitutive immunity can improve host health by lowering parasite burden while maintaining higher energy for other metabolic work.

Despite similarities in loop structure, alternate stable states emerge in IGP but not PIE models. The reason for this discrepancy hinges on production of the enemies. In principle, when each prey / parasite *“starves the enemy”* enough, the resulting positive feedback through the three species might trigger alternative stable states (alt SS; Fig. 2). In fact, IGP with a semi-chemostat like resource produces two types of them (Verdy & Amarasekare 2010). In the simpler, more typical one, either the prey or the predator dominates. In the other less common one, predator and prey coexist or the prey is excluded (Fig. 4A). If they existed, such alt SS in PIE models might explain why large infectious doses overwhelm immune clearance, leading to infection (yielding stable *E-N_1_* or *E-N_1_-I* states), while small doses can be cleared by the immune system (in a stable *E-I* state; as seen in experiments, e.g., Merrill & Cáceres 2018). Yet, we found no such alt SS in PIE or PIEc as formulated (nor did it arise in a similar model with another representation of immune functioning: Greenspoon *et al*. 2018). Hence, some other mechanism is needed to generate alt SS within hosts (e.g., effects of parasites on within-host resource supply points: van Leeuwan *et al*. 2019). In the future, such mechanisms could be added to the PIE/PIEc framework. However, they would likely break the straightforward food web analogues that we strove for here.

### Coinfection dynamics

In contrast, KP and 2PIE models produced either coexistence or priority effects of competitors for similar reasons. At first glance, such parallels might seem surprising since KP more simply connects species engaged in resource and apparent competition. In contrast, 2PIE has more complex loop structure (created by those omnivory-like *I-E* connections). Despite these structural differences, both models share qualitatively identical four species loops that govern stability (Fig. 5). We translated the net sum of those positive and negative loops into differences in *effects on* and *affected by* ratios for each competitor. Those ratios then lay out conditions for successful coinfection *vs.* alternative stable states (priority effects). First, either case requires a sufficient tradeoff. If one species competes superiorly for host energy, the other must better resist immune attack. Such a tradeoff anchors a directionality of the *affected by* ratio (immune to energy) between competing parasites. Coinfection (and coexistence more broadly), then, requires a symmetry: the parasite with greater *affected by* ratio must also have the greater *effects on* ratio (Fig. 5). Those conditions arise when the superior energy competitor without immune cells (i.e., the one with lower minimal energy needs) becomes the superior apparent competitor with them (i.e., the one that supports and withstands the highest immune density). Meanwhile, the more resistant parasite becomes the best energy competitor. If true, each parasite indirectly inhibits its own growth and facilitates its competitor. Coinfection ensues due to net negative feedback. In contrast, if one parasite always competes superiorly for energy while the superior apparent competitor enjoys resistance, asymmetry in these ratios ensues. That asymmetry means each competitor indirectly facilitates its own growth and slows its competitor, either by *“feeding the immune cells”* that attack the competitor (freeing up energy), or by *“starving the immune cells”* via starving the competitor (Fig. 5). When strong enough, such interactions ensure priority effects via net positive feedbacks.

These new within-host competition models can also guide future coinfection experiments. First, the competition models focus attention on traits and mechanism. Traditional coinfection experiments alter initial densities / order of arrival of parasites within a host (Lohr *et al* 2010; Natsopoulou *et al* 2015; *reviewed in* Karvonen *et al* 2019). Then, mechanism is inferred from pattern. Alternatively, with 2PIE-like models in hand now, experimenters can directly measure and/or fit key host-parasite traits that produce within-host dynamics. Second, they highlight how multiple niche dimensions govern competitive outcomes within hosts. Some recent empirical studies emphasized immune-(Ezenwa *et al* 2010; Halliday *et al* 2018) or resource-mediated (Budischak *et al* 2018) competition within hosts. Yet, competition along a single niche dimension predicts only competitive exclusion (*sensu stricto*). Instead, measurements of both immune and energetic niche dimensions together (like in Budischak *et al* 2015) could evaluate the entire range of outcomes anticipated by 2PIE. Third, the bifurcation maps here suggest joint manipulation of resource supply to hosts and parasite traits, perhaps leveraging genetic variation in traits among parasite strains. Such manipulations could then capture a whole array of within-host competitive outcomes (Budischak *et al* 2018) or explain increasing parasite burden along a resource gradient (Wale *et al* 2017). Finally, our model predicts that allocation to constitutive immunity strengthens indirect effects, squeezing parameter space allowing coinfection and reducing parasite burden. One could test such predictions using strains with immune knockouts (like in Chen *et al* 2020) or host genotypes differing in allocation (Fuess *et al* 2020).

## Future directions and conclusion

This within-host parasite framework could become expanded in the future. First, models could add niche dimensions. For instance, parasites might compete for multiple within-host resources (Budishack *et al* 2015; Ezenwa 2021) and/or face multiple immune defenses (Jolles *et al* 2008; Ezenwa & Jolles 2011). These additional dimensions parallel food webs with multiple resources and predators (necessitating statements of clear assumptions about specialist vs. generalist consumption, different possible tradeoffs, etc.: Hulot & Loreau 2006). These niche dimensions might expand conditions promoting coinfection or priority effects (Graham 2008; Rynkiewicz *et al* 2015). Second, other mechanisms like relative non-linearities (Armstrong & McGehee 1980), competitive intransitivity (May & Leonard 1975), or other variation-based mechanisms (Chesson 2000) may predict successful coexistence in our within-host framework. For example, since PIE can oscillate, two parasites might coexist via oscillations (relative non-linearities). Similarly, multiple parasites competing intransitively for two or more resources might coexist (Huisman & Weissing 2001). Finally, environmental variation could alter key within-host traits (*e.g.,* temperature fluctuations can modulate host immunity and parasite attack rates [Scharsack *et al* 2016]). Such intersections of environment with trait plasticity might then alter or enhance opportunities for parasite coexistence (Chesson 2000). Finally, the direct (DE) vs. indirect effect (IE) approach to predicting oscillations may have limits. Perhaps theoreticians can extend it beyond three to four or more dimensions. Together, these expansions would extend mechanistic models of within-host dynamics and produce insight into disease and coexistence alike.

In this study, we gleaned insights for within-host parasite dynamics using food web modules and feedback loops. Feedback loops enabled comparison of stability outcomes in structurally similar but biological distinct systems. For instance, similarity in loop structure in IGP and PIE successfully predicted single prey / parasite persistence either stably or via oscillations. Yet, despite these structural similarities, differences in generation of enemies (predator vs. immune cells) prevented IGP’s two alternate stable states from arising in PIE. In contrast, despite omnivory-like *I-E* connections in 2PIE, competing prey or parasites coexist (coinfect) or show priority effects like in KP. Those outcomes arose because competitors within hosts or in food webs engage in structurally similar resource and apparent competition. Such comparable structure meant that symmetries in *effects on* and *affected by* ratios (involving the enemy and resource) determined stability of competition. Hence, dynamical forces governing stability in food-webs can mirror those within hosts. Such insights can guide more predictive experiments at the within-host scale but also better population-level models. For instance, fluctuation in food supply could shift competitive outcomes within hosts that then shifts multi-parasite outbreaks at the population scale. Hence, further development of resource and immune-explicit frameworks can only enhance predictive insight into host health and disease outbreaks alike.

## ACKNOWLEDGMENTS

AR was supported by the Department of Biology, Indiana University – Bloomington. The study was supported in part by NSF DEB (grant 1655656). We thank F. Bashey-Visser, C.M Lively, and members of the Hall lab for their valuable feedback.

## Box 1.

### A feedback loop approach to analyze a general 3-dimensional system of equations

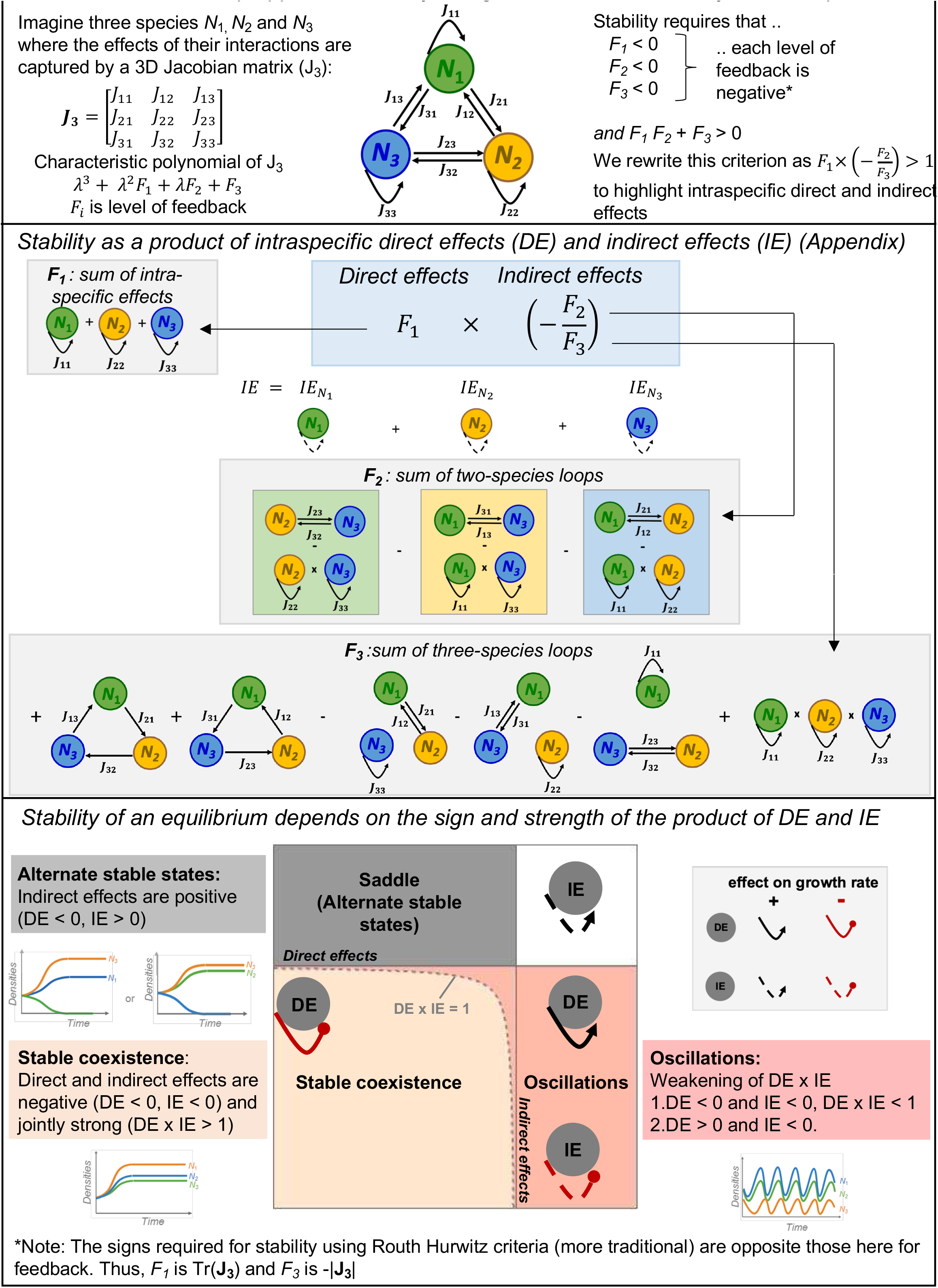

## REFERENCES

1. Alizon, S., & van Baalen, M. (2008). Multiple infections, immune dynamics, and the evolution of virulence. The American Naturalist, 172(4), E150–E168.

2. Armstrong, R. A., & McGehee, R. (1980). Competitive exclusion. The American Naturalist, 115(2), 151–170.

3. Bashey, F. (2015). Within-host competitive interactions as a mechanism for the maintenance of parasite diversity. Philosophical Transactions of the Royal Society B: Biological Sciences, 370(1675), 20140301.

4. Budischak, S. A., Wiria, A. E., Hamid, F., Wammes, L. J., Kaisar, M. M., van Lieshout, L., … & Graham, A. L. (2018). Competing for blood: the ecology of parasite resource competition in human malaria–helminth co-infections. Ecology letters, 21(4), 536–545.

5. Budischak, S. A., Sakamoto, K., Megow, L. C., Cummings, K. R., Urban Jr, J. F., & Ezenwa, V. O. (2015). Resource limitation alters the consequences of co-infection for both hosts and parasites. International Journal for Parasitology, 45(7), 455–463.

6. Chesson, P. (2000). Mechanisms of maintenance of species diversity. Annual review of Ecology and Systematics, 31(1), 343–366.

7. Chen, C. C., Louie, S., McCormick, B., Walker, W. A., & Shi, H. N. (2005). Concurrent infection with an intestinal helminth parasite impairs host resistance to enteric Citrobacter rodentium and enhances Citrobacter-induced colitis in mice. Infection and immunity, 73(9), 5468–5481.

8. Childs, L. M., & Buckee, C. O. (2015). Dissecting the determinants of malaria chronicity: why within-host models struggle to reproduce infection dynamics. Journal of The Royal Society Interface, 12(104), 20141379.

9. Clay, P. A., Cortez, M. H., Duffy, M. A., & Rudolf, V. H. (2019). Priority effects within coinfected hosts can drive unexpected population-scale patterns of parasite prevalence. Oikos, 128(4), 571–583.

10. Costantini, D., & Møller, A. P. (2009). Does immune response cause oxidative stress in birds? A meta-analysis. Comparative Biochemistry and Physiology Part A: Molecular & Integrative Physiology, 153(3), 339–344.

11. Cressler, C. E., Nelson, W. A., Day, T., & McCauley, E. (2014). Disentangling the interaction among host resources, the immune system and pathogens. Ecology letters, 17(3), 284–293.

12. De Roode, J. C., Helinski, M. E., Anwar, M. A., & Read, A. F. (2005). Dynamics of multiple infection and within-host competition in genetically diverse malaria infections. The American Naturalist, 166(5), 531–542.

13. Doublet, V., Natsopoulou, M. E., Zschiesche, L., & Paxton, R. J. (2015). Within-host competition among the honey bees pathogens Nosema ceranae and Deformed wing virus is asymmetric and to the disadvantage of the virus. Journal of invertebrate pathology, 124, 31–34.

14. Duneau, D., Ferdy, J. B., Revah, J., Kondolf, H., Ortiz, G. A., Lazzaro, B. P., & Buchon, N. (2017). Stochastic variation in the initial phase of bacterial infection predicts the probability of survival in D. melanogaster. Elife, 6, e28298.

15. Ezenwa, V. O. (2021). Co-infection and nutrition: integrating ecological and epidemiological perspectives. In Nutrition and Infectious Diseases (pp. 411–428). Humana, Cham.

16. Ezenwa, V. O., & Jolles, A. E. (2011). From host immunity to pathogen invasion: the effects of helminth coinfection on the dynamics of microparasites. Integrative and comparative biology, 51(4), 540–551.

17. Ezenwa, V. O., Etienne, R. S., Luikart, G., Beja-Pereira, A., & Jolles, A. E. (2010). Hidden consequences of living in a wormy world: nematode-induced immune suppression facilitates tuberculosis invasion in African buffalo. The American Naturalist, 176(5), 613–624.

18. Fenton, A., & Perkins, S. E. (2010). Applying predator-prey theory to modelling immune-mediated, within-host interspecific parasite interactions. Parasitology, 137(6), 1027–1038.

19. Fuess, L. E., Weber, J. N., den Haan, S., Steinel, N. C., Shim, K. C., & Bolnick, D. I. (2020). A test of the Baldwin Effect: Differences in both constitutive expression and inducible responses to parasites underlie variation in host response to a parasite. bioRxiv.

20. Gorsich, E. E., Etienne, R. S., Medlock, J., Beechler, B. R., Spaan, J. M., Spaan, R. S., … & Jolles, A. E. (2018). Opposite outcomes of coinfection at individual and population scales. Proceedings of the National Academy of Sciences, 115(29), 7545–7550.

21. Graham, A. L. (2008). Ecological rules governing helminth–microparasite coinfection. Proceedings of the National Academy of Sciences, 105(2), 566–570.

22. Greenspoon, P. B., Banton, S., & Mideo, N. (2018). Immune system handling time may alter the outcome of competition between pathogens and the immune system. Journal of theoretical biology, 447, 25–31.

23. Griffiths, E. C., Fairlie-Clarke, K., Allen, J. E., Metcalf, C. J. E., & Graham, A. L. (2015). Bottom-up regulation of malaria population dynamics in mice co-infected with lung-migratory nematodes. Ecology letters, 18(12), 1387–1396.

24. Halliday, F. W., Umbanhowar, J., & Mitchell, C. E. (2018). A host immune hormone modifies parasite species interactions and epidemics: insights from a field manipulation. Proceedings of the Royal Society B, 285(1890), 20182075.

25. Hite, J. L., & Cressler, C. E. (2018). Resource-driven changes to host population stability alter the evolution of virulence and transmission. Philosophical Transactions of the Royal Society B: Biological Sciences, 373(1745), 20170087.

26. Holt, R. D., & Polis, G. A. (1997). A theoretical framework for intraguild predation. The American Naturalist, 149(4), 745–764.

27. Holt, R. D., Grover, J., & Tilman, D. (1994). Simple rules for interspecific dominance in systems with exploitative and apparent competition. The American Naturalist, 144(5), 741–771.

28. Hulot, F. D., & Loreau, M. (2006). Nutrient-limited food webs with up to three trophic levels: feasibility, stability, assembly rules, and effects of nutrient enrichment. Theoretical Population Biology, 69(1), 48–66.

29. Hurwitz, A. (1895). On the conditions under which an equation has only roots with negative real parts. Math. Ann. 46, 273–284 (1895)

30. Jolles, A. E., Ezenwa, V. O., Etienne, R. S., Turner, W. C., & Olff, H. (2008). Interactions between macroparasites and microparasites drive infection patterns in free-ranging African buffalo. Ecology, 89(8), 2239–2250.

31. Huisman, J., & Weissing, F. J. (2001). Fundamental unpredictability in multispecies competition. The American Naturalist, 157(5), 488–494.

32. Karvonen, A., Jokela, J., & Laine, A. L. (2019). Importance of sequence and timing in parasite coinfections. Trends in parasitology, 35(2), 109–118.

33. Leibold, M. A. (1996). A graphical model of keystone predators in food webs: trophic regulation of abundance, incidence, and diversity patterns in communities. The American Naturalist, 147(5), 784–812.

34. Lohr, J. N., Yin, M., & Wolinska, J. (2010). Prior residency does not always pay off–co-infections in Daphnia. Parasitology.

35. May, R. M., & Leonard, W. J. (1975). Nonlinear aspects of competition between three species. SIAM journal on applied mathematics, 29(2), 243–253.

36. Merrill, T. E. S., & Cáceres, C. E. (2018). Within-host complexity of a plankton-parasite interaction. Ecology, 99(12), 2864–2867.

37. Metcalf, C. J. E., Grenfell, B. T., & Graham, A. L. (2020). Disentangling the dynamical underpinnings of differences in SARS-CoV-2 pathology using within-host ecological models. PLoS pathogens, 16(12), e1009105.

38. Mideo, N., Alizon, S., & Day, T. (2008). Linking within-and between-host dynamics in the evolutionary epidemiology of infectious diseases. Trends in ecology & evolution, 23(9), 511–517.

39. Natsopoulou, M. E., McMahon, D. P., Doublet, V., Bryden, J., & Paxton, R. J. (2015). Interspecific competition in honeybee intracellular gut parasites is asymmetric and favours the spread of an emerging infectious disease. Proceedings of the Royal Society B: Biological Sciences, 282(1798), 20141896.

40. Novak, M., Yeakel, J. D., Noble, A. E., Doak, D. F., Emmerson, M., Estes, J. A., … & Wootton, J. T. (2016). Characterizing species interactions to understand press perturbations: what is the community matrix?. Annual Review of Ecology, Evolution, and Systematics, 47, 409–432.

41. Otterstatter, M. C., & Thomson, J. D. (2006). Within-host dynamics of an intestinal pathogen of bumble bees. Parasitology, 133(6), 749.

42. Puccia, C. J., & Levins, R. (1991). Qualitative modeling in ecology: loop analysis, signed digraphs, and time averaging. In Qualitative simulation modeling and analysis (pp. 119–143). Springer, New York, NY.

43. Råberg, L. (2012). Infection intensity and infectivity of the tick-borne pathogen Borrelia afzelii. Journal of evolutionary biology, 25(7), 1448–1453.

44. Råberg, L., Graham, A. L., & Read, A. F. (2009). Decomposing health: tolerance and resistance to parasites in animals. Philosophical Transactions of the Royal Society B: Biological Sciences, 364(1513), 37–49.

45. Routh, E. J. (1877). A Treatise on the Stability of a Given State of Motion, Particularly Steady Motion: Being the Essay to which the Adams Prize was Adjudged in 1877, in the University of Cambridge. Macmillan and Company.

46. Rynkiewicz, E. C., Pedersen, A. B., & Fenton, A. (2015). An ecosystem approach to understanding and managing within-host parasite community dynamics. Trends in parasitology, 31(5), 212–221.

47. Scharsack, J. P., Franke, F., Erin, N. I., Kuske, A., Büscher, J., Stolz, H., … & Kalbe, M. (2016). Effects of environmental variation on host–parasite interaction in three-spined sticklebacks (Gasterosteus aculeatus). Zoology, 119(4), 375–383.

48. Susi, H., Barrès, B., Vale, P. F., & Laine, A. L. (2015). Co-infection alters population dynamics of infectious disease. Nature communications, 6(1), 1–8.

49. Van Leeuwen, A., Budischak, S. A., Graham, A. L., & Cressler, C. E. (2019). Parasite resource manipulation drives bimodal variation in infection duration. Proceedings of the Royal Society B, 286(1902), 20190456.

50. Verdy, A., & Amarasekare, P. (2010). Alternative stable states in communities with intraguild predation. Journal of Theoretical Biology, 262(1), 116–128.

51. Wodarz, D. (2006). Ecological and evolutionary principles in immunology. Ecology Letters, 9(6), 694–705.

52. Wolfram Research, Inc., Mathematica, Version 12.2, Champaign, IL (2020)

