## Supplemental Appendix for "Niche theory for within-host parasite dynamics: Analogies to food web modules via feedback loops"

In this supplemental appendix, we provide mathematical support to underlie the bifurcation diagrams and loop approaches in the main text. Section 1 describes the general approach to stability and loop analysis in three dimensions (Box 1). Section 2 works through the three dimension models of an omnivorous food web (intraguild predation, IGP) and its within-host analogues (PIE and PIEc; Table 1; Figs. 2-4). In section 3, we provide a similar analysis for the model of joint apparent and resource competition in the food web (keystone predation, KP) and its within-host equivalent (2PIE; Table 2; Figs. 5,6). All analyses used Mathematica Version 12 (Wolfram Research Inc., Champaign, IL. USA).

### Section 1: Calculating intraspecific direct (DE) and indirect effects (IE) in three dimensions using Jacobian and inverse Jacobian matrices – a general approach

Imagine we have species $N_1$, $N_2$, and $N_3$ interacting according to differential equations $dN_1/dt = G_1$, $dN_2/dt= G_2$, and $dN_3/dt = G_3$ (and $G_j$ are growth rate functions). A 3D Jacobian ($\mathbf{J_3}$) matrix captures the intra- and inter-specific direct effects of these interactions (as $\partial G_i/\partial N_j$) while the (negative) elements of the inverse Jacobian matrix ($-\mathbf{J_3}^{-1}$) captures the intra- and inter-specific indirect effects of these interactions (Box 1). The Jacobian for this three-dimensional system is:

$$\boldsymbol{J}_3 = \begin{bmatrix} J_{11} & J_{12} & J_{13} \\ J_{21} & J_{22} & J_{23} \\ J_{31} & J_{32} & J_{33} \end{bmatrix} \quad \text{(A1)}$$

where intraspecific direct effects of species $i$, $J_{ii}$, fall along the main diagonal, and inter-specific effect of $j$ on $i$ fall in the $J_{ij}$ off-diagonal elements. Such a three species system then has three corresponding levels of feedback ($F_i$):

Level 1, $F_1 = J_{11} + J_{22} + J_{33}$ (A2.a)

Level 2, $F_2 = (J_{23} J_{32} - J_{22} J_{33}) + (J_{13} J_{31} - J_{11} J_{33}) + (J_{12} J_{21} - J_{11} J_{22})$ (A2.b)

Level 3, $F_3 = J_{13} J_{21} J_{32} + J_{12} J_{31} J_{23} -$ (A2.c)

$(J_{12} J_{21} J_{33} + J_{13} J_{31} J_{22} + J_{23} J_{32} J_{11}) + J_{11} J_{22} J_{33}$

where $F_1$ (equ. A2.1) is the trace of the Jacobian, tr[$\mathbf{J_3}$], and is the sum of intraspecific direct effects. Level two feedback, $F_2$ (equ. A2.2), is the sum of two species interactions, and $F_3$ (equ. A2.c) is the sum of three species lops. The inverse Jacobian, $\mathbf{J_3}^{-1}$, then, captures those intraspecific indirect effects on the main diagonal; it is:

$$\boldsymbol{J_3}^{-1} = \begin{bmatrix} \frac{J_{23}J_{32}-J_{22}J_{33}}{F_3} & \frac{J_{12}J_{32}-J_{13}J_{32}}{F_3} & \frac{J_{13}J_{22}-J_{12}J_{23}}{F_3} \\ \frac{J_{21}J_{33}-J_{23}J_{31}}{F_3} & \frac{J_{13}J_{31}-J_{11}J_{33}}{F_3} & \frac{J_{11}J_{23}-J_{13}J_{21}}{F_3} \\ \frac{J_{22}J_{31}-J_{21}J_{32}}{F_3} & \frac{J_{11}J_{32}-J_{12}J_{31}}{F_3} & \frac{J_{12}J_{21}-J_{11}J_{22}}{F_3} \end{bmatrix} \quad \text{(A3)}$$

Specifically, the negative of the elements of the inverse Jacobian matrix, $\mathbf{-J_3^{-1}}$, $-J^{-1}_{ij}$, show the net effect of species $j$ on species $i$ due the direct linkage with species $i$ and all possible indirect pathways through which species $i$ and $j$ are connected via linkages with intermediate species (based on Novak *et al* 2016). Notice, then, how the elements falling on the main diagonal correspond to the intraspecific indirect effects described in the text, where:

$IE_{N_1}$: The indirect effect of $N_1$ on itself, $J_{11}^{-1}$: $-(J_{23} J_{32} - J_{22} J_{33}) / F_3$ (A4.a)

$IE_{N_2}$: The indirect effect of $N_2$ on itself, $J_{22}^{-1}$: $-(J_{13} J_{31} - J_{11} J_{33}) / F_3$ (A4.b)

$IE_{N_3}$: The indirect effect of $N_3$ on itself, $J_{33}^{-1}$: $-(J_{12} J_{21} - J_{11} J_{22}) / F_3$ (A4.c)

where $F_3$ follows above (eq. A2.c). Furthermore, the numerators of these three *IE* terms add up to level two feedback, $-F_2$ (eq. A2.b). Hence, the sum of the indirect effects of each species on itself (the trace of $\mathbf{-J_3^{-1}}$) is $-F_2 / F_3$. (Below, we show a related result for the four dimension models). Stability requires that the product of intraspecific direct ($F_1$) and indirect effects ($- F_2 / F_3$) exceeds 1, or $F_1 ( - F_2 / F_3 ) >$ which is equivalent to saying that *DE* x *IE* > 1 (Box 1).

**Section 2: Analysis of the three dimension models (IGP, PIE, and PIE$_C$)**

In this section, we provide analysis of the three dimension models, especially that of IGP (Table 1, Figs. 2-4). Then, we detail some aspects of the PIE and PIE$_C$ models. For simplicity in this section, we drop the '1' notation, so $N_1$ is simply $N$.

*System of equations (eq. 1, 3; see also Table 1)*

*The growth rate of the top consumer,* P *or* I: In IGP (Eq. 1A), the predator (*P*) feeds on resources (*R*) at rate $f_{PR}$ and on prey ($N_1$) at rate $f_{PN_1}$. Predators convert consumed prey into births with conversion $e_{PN_1}$ and die at background rate $m_P$. In PIE (Eq. 2A), immune cells (*I*) attack parasite ($N_1$) at rate $f_{IN_1}$. Immune cells increase when that interaction induces consumption of host energy, *E*, with per parasite proportion $e_{IN_1}$. They also increase in PIEc as *E* is allocated at baseline rate $a_b$. The energy – induced or allocated – is converted into immune cells with efficiency $e_I$. Finally, immune cells are lost at rate $m_I$.

*The growth rate of the intermediate consumer*, *prey or parasite* $N_1$: In IGP (Eq. 1B), prey feed on resources at rate $f_{N_1}$ that convert basal resources into births. Similarly, in PIE (Eq. 2B), parasites feed on energy at rate $f_{N_1}$and with conversion efficiency $e_{N_1E}{}^{-1}$. The shared symbol for feeding rate of prey or parasite ($N_1$) facilitates comparison across models. In both models, the prey/ parasite are lost due to attack by the top consumer ($f_{PN_1}$/ $f_{IN_1}$) and die at background rate $m_N$.

*The growth rate of the resource*, R *or* E: In IGP (Eq. 1C), the resource follows chemostat renewal, with supply point *S* and loss (dilution) rate *a*. It is then consumed by prey ($N_1$) and predators (*P*) having stoichiometric conversions $Q_{N_1}$and $Q_P$,respectively. In the PIE (Eq. 2C), energy is supplied similarly. The host consumes resource *S* via a Monod function with maximal assimilation rate $f_E$ and half-saturation constant *h.* (This function merely pays homage to nonlinear feeding behavior of hosts). That resource, converted to energy within the host (*E*), is lost at fixed rate *r* (analogous to *a*) for use by hosts (for metabolic needs). Hence, both *R* and *E* follow a donor-control type dynamic (*i.e.,* production of both decreases with density; Murdoch *et al* 2003). Additionally, host energy is consumed via induction from immune attack on parasites (proportional to the triple product $N_1 E I$), consumption (theft) by parasites, and, in PIEc, baseline allocation to immune function (at rate $a_b$).

(A) Intra-guild predation (IGP)

We consider the simplest possible model of IGP – a chemostat-like resource, *R*, fed upon by two consumers, the IGP prey (*N*) and IGP predators (*P*). Both species have linear functional responses (eq. 1, Table 1) that govern the underlying consumer-resource interactions.

*The Jacobian matrix and feedback loops*

The Jacobian, evaluated at a feasible interior (*int*) equilibrium, is (in order of *R*, *N*, *P*):

$$\mathbf{J_{IGP}} = \begin{bmatrix} F_1 & -f_N Q_N R^*_{int} & -f_N Q_N R^*_{int} \\ f_N N^*_{int} & 0 & -f_{PN} N^*_{int} \\ f_{PR} P^*_{int} & e_{PN} f_{PN} P^*_{int} & 0 \end{bmatrix} \qquad \text{(A5)}$$

where $F_1$ is level 1 feedback and the self-limitation of the resource, $J_{RR}$, (also Eq. 6.a) and level 2 and 3 feedback, $F_2$ and $F_3$, are (after eq. A2.b,c respectively):

$$F_1 = J_{RR} = -a - f_N Q_N N_{int}^* - f_{PR} Q_P P_{int}^* = -a S / R_{int}^* < 0 \qquad \text{(A6.a)}$$

$$F_2 = J_{RN} J_{NR} + J_{PN} J_{NP} + J_{RP} J_{PR} =$$

$$-e_{PN} f_{PN}^2 N_{int}^* P_{int}^* - ( f_N^2 Q_N N_{int}^* + f_{PR}^2 Q_P P_{int}^*) R_{int}^* < 0. \qquad \text{(A6.b)}$$

$$F_3 = (i) + (ii) + (iii) = (J_{PN} J_{NP} J_{RR}) + (J_{PN} J_{RP} J_{NR}) + (J_{RN} J_{PR} J_{NP}) \qquad \text{(A6.c)}$$

Both $F_1$ and $F_2$ are always negative. $F_2$ is also always negative because it is the sum of three pairwise two-species consumer-resource [+/-] loops – prey resource, predator-prey, and predator-resource, respectively (eq. A6.b). Then, $F_3$ is the sum of three loops – (i) '$N_1$ *is eaten*', (ii) '*starving the enemy*', and (iii) '*feeding the enemy*' (Fig. 2). We describe $F_3$ in more detail below. Unfortunately, the condition for oscillations for IGP (and the PIE models below), $F_1 F_2 + F_3 < 0$, or $F_1 (-F_2 / F_3) < 1$ (Box 1), is not analytically tractable (but readily conquered with a computer).

*Building the bifurcation diagram*

We can then build the bifurcation diagrams. We created them along gradients of feeding rate and supply point ($f_N$-$S$). We focus on details of the version with alternative stable states (Fig. 4A) shown in more detail here (Fig. A1.A). First, the IG prey $N$ can invade the $R$-only system and reach its $R_N^*$-$N_R^*$ equilibrium with sufficient nutrient supply, when $S > R_N^*$; similarly, IG predators can invade the $R$-only system and reach its $R_P^*$-$P_R^*$ equilibrium when $S > R_P^*$, where:

$$R_N^* = \frac{m_N}{f_N} \quad \text{(A7.a)}$$

$$N_R^* = \frac{a(S-R_N^*)}{Q_N f_N} \quad \text{(A7.b)}$$

$$R_P^* = \frac{m_P}{f_{PR}} \quad \text{(A7.c)}$$

$$P_R^* = \frac{a(S-R_P^*)}{Q_P f_{PR}} \quad \text{(A7.d)}$$

and where minimal resource requirements ($R_N^*$, $R_P^*$) are ratios of mortality to birth rate per unit resource (after stoichiometric conversion), and consumer densities ($N_R^*$, $N_P^*$) are ratios of resource production to per capita consumption (with conversion). We imagine that a tradeoff could permit coexistence: hence, the IG prey is the superior resource competitor ($R_N^* < R_P^*$; their competition becomes more asymmetric as $R_P^*$ becomes relatively larger; $f_N$ must exceed the intersection of the four curves in Fig. A1.A). We can also define a recurrent $N_P^*$, the density of

IG prey needed to support if it only ate prey (i.e., it was not omnivorous):

$$N_P^* = \frac{m_P}{e_{PN} f_{PN}} \quad \text{(A8)}$$

Then, from equations for the IGP prey ($dN/dt$) and IG predator ($dR/dt$), we can also determine that a feasible interior (*int*) requires an intermediate $R_{int}^*$ value, one which falls between those minimal resource requirements of the consumers alone, $R_N^* < R_{int}^* < R_P^*$:

from $dN/dt$ (eq. 1b): $P_{int}^* = f_N (R_{int}^* - R_N^*) > 0$, so $R_N^* < R_{int}^*$ (A9.a)

from $dP/dt$ (eq. 1c): $N_{int}^* = N_P^* ( R_P^* - R_{int}^*) / R_P^* > 0$, so $R_{int}^* < R_P^*$ (A9.b)

Otherwise, one species excludes the other (e.g., in the *R-N* region of Fig. A1.A, $R_{int}^* > R_P^*$; $R_{int}^*$ is not real in the *R-P* region below the fold [*F*, see below]).

The IG prey, *N*, can invade the *R-P* system when there is enough resource, *R*, to offset mortality from the IG predator, *P*:

$$f_N (R_P^* - R_N^*) > f_{PN} P_R^* \quad \text{(A10.a)}$$

$$\text{or } S < R_P^* + \frac{f_N}{a}\left(1 - \frac{R_N^*}{R_P^*}\right)(f_{PR} Q_P R_P^*) = T_2 \quad \text{(A10.b)}$$

which requires that *S* fall between $R_P^*$ and $T_2$ (i.e., to the left of $T_2$ on Fig. A1.A). In turn, the IG predator can invade the *R-N* system when there is enough prey at the *R-N* boundary, $N_R^*$ (eq. A7.b) to offset its competitive disadvantage (yet still less than $N_P^*$):

$$N_R^* > N_P^* \left(1 - \frac{R_N^*}{R_P^*}\right) \quad \text{(A11.a)}$$

$$\text{or } S > R_N^* + \frac{f_N}{a}\left(1 - \frac{R_N^*}{R_P^*}\right)(N_P^*) = T_1 \quad \text{(A11.b)}$$

(i.e. to the right of $T_1$ on Fig. A1.A). This threshold $T_1$ reaches a vertical asymptote at:

$$S = \frac{f_N R_N^* N_P^* Q_N}{a} \quad \text{(A12)}$$

which happens when flow of resources in to the system, *a S*, equals the consumption of resources by the minimal density of prey needed to support the predators, $f_N R_N^* N_P^* Q_N$. When *S* exceeds this threshold (eq. A13), the prey can longer exclude the predator (but the predator could still exclude the prey).

The two boundary equilibria are mutually invasible when:

$$aS > f_N R_N^* N_P^* (Q_N - e_{PN} Q_P) \quad \text{(A13)}$$

(which sits above where $T_1$ and $T_2$ cross the second time along an $S$ gradient on Fig. A1.A). Notice how mutual invasion would be assured if

$$Q_N < e_{PN} Q_P \quad \text{(A14)}$$

i.e., if the IG prey is more efficient than the joint efficiency of the IG predator. This trait arrangement guarantees that the sum of the '*starving the enemy*' (loop ii, negative; Fig. 2) and '*feeding the enemy*' (loop iii, positive) components of level 3 feedback, $F_3$, is negative since:

loop (ii): $J_{PN}\, J_{RP}\, J_{NR} = -\, e_{PN} f_N f_{PN} f_{PR}\, Q_P\, N^*\, P^*\, R^*$ (A15.a)

loop (iii): $J_{RN}\, J_{PR}\, J_{NP} = f_N f_{PN} f_{PR}\, Q_N\, N^*\, P^*\, R^*$ (A15.b)

loops (ii) + (iii) $= f_N f_{PN} f_{PR}\, N^*\, P^*\, R^*\, (Q_N - e_{PN}\, Q_P)$ (A15.c)

If those two loops are net negative – even without the stabilizing third '*N is eaten*' (loop [i], Fig. 2) - then the interior always yields coexistence when feasible (as in Fig. 3). Conversely, when $Q_N > e_{PN}\, Q_P$, other possibilities arise (Figs. 4, A1). Specifically, the *R-N* and *R-P* boundaries are mutually uninvasible between this intersection (eq. A14, but above $S = R_P^*$, where $T_1 = T_2$ again). Hence, unlike in keystone predation (section 3), mutual invasion is a phenomenon arising at higher supply (S), not lower. Assuming that $Q_N > e_{PN} Q_P$, a fold bifurcation (*F* in Fig. A1.A) becomes possible; it sits at:

$$S_F = \frac{\left(a + f_N (N_P^* Q_N - f_{PR} R_N^* Q_P)\right)^2}{4 a f_N (N_P^*/R_P^*)(Q_N - e_{PN} Q_P)} \quad \text{(A16)}$$

When $S$ falls below this threshold ($S < S_F$; left of line *F* in Fig. A1.A), two interior equilibria become feasible. Above it, both interior equilibria vanish (i.e., become complex numbers, leaving only a stable *R-P* boundary). This fold bifurcation intersects with $T_1$ (dot: Fig. A1.A) at:

$$S_{T-F} = \frac{f_N R_N^* N_P^* Q_N (Q_N - e_{PN} Q_P)}{a\left((2Q_N - e_{PN} Q_P) - a/\left(f_{PR} R_N^*\right)\right)} \quad \text{(A17)}$$

an expression which defies transparent explanation (at least to us). However, the 'Alt SS-2' region requires that $S < S_{T\text{-}F}$. Assuming that is true, stability of one of the two interior equilibria is possible when its $F_3 < 0$; the other is a saddle ($F_3 > 0$). This third level of feedback, when all three loops are added together, (i) + (ii) + (iii):

$$F_3 = f_{PN} N^* P^* \left( f_N f_{PR} R_P^* / N_P^* (Q_N - e_{PN} Q_P) + e_{PN} f_{PN} F_1 \right) \quad \text{(A18)}$$

is negative when:

$$aS < f_N (R_P^* / N_P^*)(Q_N - e_{PN} Q_P) {R_{int}^*}^2 \quad \text{(A19)}$$

When the fold is present, this condition enables one of the feasible interior equilibria to be stable in the region bound by $T_2$, $T_1$, and *F* (Alt SS-2; Fig. A1.A). Notice that this region sits outside that bounding mutual invasion (between $T_1$ and $T_2$; eq. A14). Hence, we are reminded that coexistence can occur in IGP in parameter space outside that granting mutual invasibility.

*Positive intraspecific indirect feedback leads to alternate stable states in IGP*

To summarize, using certain parameter values (that allow $Q_N > e_{PN} Q_P$; Fig. A1.A), a 2D bifurcation diagram of IGP shows six different regions in the space of supply point (*S*) and feeding rate of the prey ($N_1$) on resource ($f_{N_1}$). There, a fold (*F* [eq. A16], for $f_{N_1}$ above dot [eq. A17]) and two transcritical ($T_1$ [eq. A10.b] and $T_2$ [eq. A11.b] bifurcations separate a resource only (*R*) equilibrium from two-species boundary equilibria (*R*-$N_1$, *R-P*) and from situations with one or two feasible interior equilibria (with $Z_1$, $Z_2$; Fig. A1.A). To understand these regions, it helps to combine summed intraspecific indirect effects (IE; Fig. A1.B,C) for each equilibrium feasible (stable or not) with changes in the equilibria themselves (Fig. A1.D, E). Taking a slice of parameter space at low supply point (*S*=24), the IGP system starts with stable *R*-*P* (IE<0) at low $f_{N_1}$; the other *R*-$N_1$ boundary is a saddle (IE>0). A fold then creates two interiors which lead to three-species coexistence ($Z_1$: IE<0) or exclusion of prey (because $Z_2$ is a saddle: IE>0; the resource-predator boundary is still stable [IE < 0]). At the first transcritical ($T_1$), the stable interior becomes infeasible but the prey boundary becomes stable, so either prey or predator persists alone (for both *R*-$N_1$ and *R*-*P*: IE <0; for $Z_2$: IE > 0 still). Past the second transcritical ($T_2$), only the prey-resource remains stable ($Z_2$ is no longer feasible; *R*-*P*: IE < 0). At higher nutrient supply (*S*=40, Fig. A1.C,E), simpler transitions occur with increasing $f_{N_1}$. First, the fold introduces the stable interior ($Z_1$) to the stable *R*-*P* boundary (separated by the saddle interior: $Z_2$). Then at $T_2$, the saddle ($Z_2$) becomes infeasible and only predator-prey coexistence is possible (IE<0).

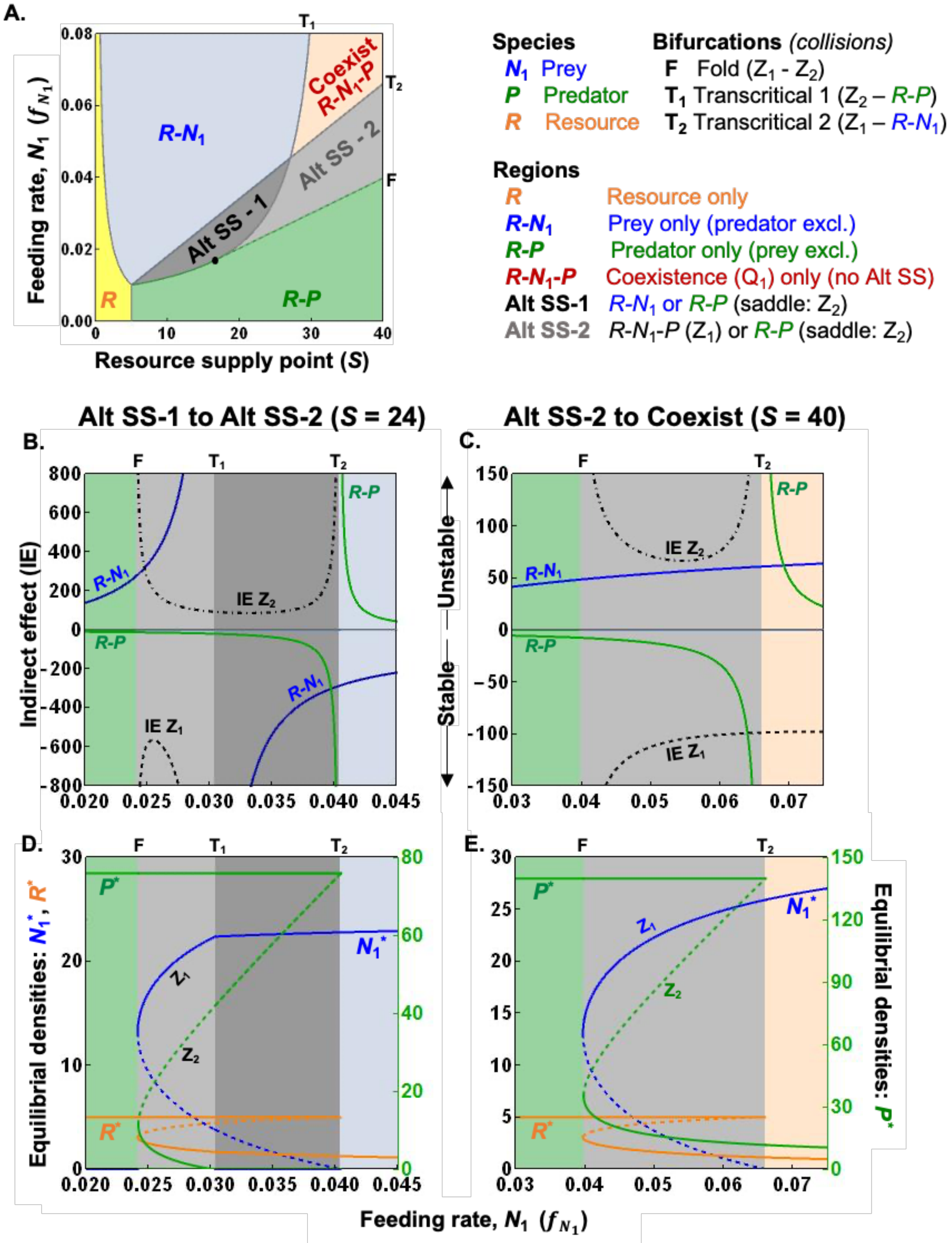


**Figure A1**: *Positive intraspecific indirect feedback leads to two forms of alternate stable states (Alt SS-1 and -2) in the IGP model of prey ($N_1$), predators (P), and resources (R).* (A) A bifurcation diagram in resource supply point (*S*) - prey feeding rate ($f_{N_1}$) space. A fold (F, for $f_{N_1}$ above dot) and transcritical ($T_1$, $T_2$ noted) bifurcations create six regions (see key) composed of boundary (*R, R-*$N_1$*, R-P*) or interior ($Z_1$, $Z_2$) equilibria. Along a gradient of $f_{N_1}$: (B,C) sum of intraspecific direct (DE) and indirect (IE) effects, and (D,E) densities of feasible equilibria (stable or saddle). (B,D, at lower nutrient supply [S=24]): At low $f_{N_1}$, the IGP system starts with stable R-P (IE<0). At the fold, the prey can invade, and then either predators and prey coexist (interior $Z_1$: IE<0) or predator exclude prey ($Z_2$ is the saddle: IE>0). Past $T_2$, either prey or predator persists (IE for *R-*$N_1$ and *R-P* both positive; $Z_2$ is still the saddle). Past $T_1$, only the prey persists ($Z_2$ not feasible). (C,E, at higher S=40): transitions occur similarly from *R-P* only until Alt SS-2 ($Z_1$ or *R-P*). At $T_1$, the saddle ($Z_2$) becomes infeasible and only coexistence is possible (IE<0). Parameter values follow defaults (Table 1).

<u>(B) One-parasite within-host model (PIE)</u>

The within-host analogue of IGP, PIE, has host energy *E*, parasites, *N* (dropping the '1' subscript here), and immune cells, *I*. Connections between *E* and *I* create similarities between PIE and the omnivorous IGP model. However, generation of immune cells from the product of energy and parasites, *EN*, changes key aspects (as described below).

*The Jacobian matrix and feedback loops*

The Jacobian for PIE (eq. 2, Table 1), evaluated at a feasible interior (*int*) equilibrium ($\mathbf{J}_{\mathbf{PIE}}$), is (in order of *E*, *N*, and *I*):

$$\mathbf{J}_{\mathbf{PIE}} = \begin{bmatrix} F_1 & -(f_N + e_{IN} f_{IN} I^*_{int}) E^*_{int} & -e_{IN} f_{IN} N^*_{int} E^*_{int} \\ f_N N^*_{int}/e_{NE} & 0 & -f_{IN} N^*_{int} \\ e_{IN} f_{IN} N^*_{int} I^*_{int}/e_I & e_{IN} f_{IN} E^*_{int} I^*_{int}/e_I & 0 \end{bmatrix} \quad \text{(A20)}$$

where feedback at levels 1-3, $F_1$ (eq. 6.b), $F_2$, and $F_3$, respectively:

$$F_1 = J_{EE} = -r - f_N N^* - e_{IN} f_{IN} I^* N^* < 0 \quad \text{(A21.a)}$$

$$F_2 = J_{EN} J_{NE} + J_{IE} J_{EI} + J_{IN} J_{NI} < 0 \quad \text{(A21.b)}$$

$$F_3 = -\frac{e_{IN} f_{IN} f_N E^* I^* N^* (e_{IN} f_N E^* N^* + e_{NE} r)}{e_I e_{NE}} < 0 \quad \text{(A21.c)}$$

are all always negative. Host energy is self-limiting ($F_{EE} < 0$, so $F_1 < 0$; eq. A21.a). As in IGP, $F_2$ is always negative just based on signs alone, since it is the sum of pairwise consumer-resource-like loops (parasite-energy, parasite-energy, and parasite-immune, respectively; eq. A21.b). Mathematically, $F_3 < 0$ (Eq. A21.c). Hence, PIE cannot produce alternative stable states – mathematically, a feasible interior cannot become a saddle because negative feedback prevails.

*Building the bifurcation diagram*

To ease notation, we define inflow of energy from the host as $f(S) = f_E S/(h+S)$. Without a parasite, energy in the host reaches the trivial equilibrium, $E_t^* = f(S) / r$ (i.e., the ratio of inputs from feeding to allocation rate to metabolic work). The parasite can invade when its minimal energy requirement, $E_N^*$, is met, yielding the boundary ($b$) equilibrium:

$$E_N^* = \frac{e_{NE} m_N}{f_N} \quad \text{(A22.a)}$$

$$N_b^* = \frac{f(S)/E_N^* - r}{f_N} \quad \text{(A22.b)}$$

$$EN_b^* = \frac{f(S) - rE_N^*}{f_N} \quad \text{(A22.c)}$$

which requires for feasibility ($N_b^* > 0$) that internal energy at the trivial equilibrium exceed the minimal energy requirement (or that $E_t^* > E_N^*$). Note that, at this boundary, energy decreases while parasite density increases with feeding rate of the parasite, $f_N$ – hence their product, $EN_b^*$ (eq. A22.c) is hump-shaped with $f_N$. Immune cells reach positive density (and an interior equilibrium becomes feasible) if that unimodal $EN_b^*$ exceeds the minimal $EN_I^*$ requirement of the parasite, where:

$$EN_I^* = \frac{e_I m_I}{e_{IN} f_{IN}}. \qquad \text{(A23)}$$

This condition is met between a lower (*L*) and higher (*H*) feeding rate of the parasite, $f_N$:

$$f_{N,L\&H} = 0.5\left(f(S)/(EN_I^*) \pm \sqrt{[f(S)/(EN_I^*)]^2 - 4e_{NE}m_N r/(EN_I^*)}\right) \qquad \text{(A24.a,b)}$$

which envelopes the stable coexistence region in the bifurcation diagram (Fig. 3B). That coexistence region, in turn, requires that the resource supply, $f(S)$ exceed a minimum to be real:

$$f(S) > 2\sqrt{e_{NE}m_N r(EN_I^*)} \qquad \text{(A25)}$$

which is where $f_{N,L} = f_{N,H}$ (from eq. A24). With this condition (equ. A25) met, feeding rate of the parasite must fall between $f_{N,L}$ and $f_{N,H}$ (equ. A24). If feeding rate is too low, the parasite is not dense enough to sustain an immune response; if it is too high, the parasite depresses energy too much (and outcompetes the immune system). When feasible, the interior equilibrium is:

$$E_{int}^* = \frac{e_{NE}(f(S)-f_N(EN_I^*))+e_{IN}f_N(EN_I^*)E_N^*}{e_{IN}f_N(EN_I^*)+e_{NE}r} \qquad \text{(A26.a)}$$

$$N_{int}^* = (EN_I^*)/E_{int}^* \qquad \text{(A26.b)}$$

$$I_{int}^* = \frac{f_N(f(S)-f_N(EN_I^*)-rE_N^*)}{e_I f_N m_I+e_{NE}r} \qquad \text{(A26.c)}$$

which requires that energy supply, $f(S)$, exceeds that consumed by the parasite at the immune system's minimal requirement, $f_N\,(EN_I^*)$ and allocated by hosts given the minimal energy requirement of the parasite, $r\,E_N^*$. (The feasibility condition for $E_{int}^* > 0$ proves less restrictive).

<u>(C) One-parasite within-host model with baseline allocation to immune cells (</u>PIE$_C$)

If the host allocates any energy to immune cells (in PIE$_C$; $a_b > 0$), immune cells are always present (i.e., there is no minimum value of $EN_I$ required for the immune system). The *E-I* boundary equilibrium now becomes:

$$E_b^* = \frac{f(S)}{a_b + r} \quad \text{(A27.a)}$$

$$I_b^* = \left(\frac{a_b}{e_I m_I}\right) E_b^* \quad \text{(A27.b)}$$

where both energy and immune system increase with influx energy from food to the host, $f(S)$. The parasite can invade this boundary (and create the *E-N-I* interior of Fig. 3C) when it has positive fitness at the boundary, i.e., when:

$$\frac{E_b^*}{E_N^*} > \frac{f_{IN} I_b^* + m_N}{m_N} \quad \text{(A28)}$$

where $E_N^*$ is a minimal energy requirement of the parasite assuming no mortality from immune cells (and $E_N^* = e_N\, m_N / f_N$). This invasion criterion first requires that $E_b^* > E_N^*$ - the energy available ($E_b^*$) must exceed that minimum without immune killing ($E_N^*$). However, that excess must compensate for the added mortality from immune cells ($f_{IN} I_b^* + m_N$) relative to background mortality ($m_N$).

Once the parasite can invade, the interior equilibrium of the PIEc model becomes rather algebraically unwieldly, unfortunately. The Jacobian, evaluated at the interior, changes, too (from eq. A20). Now, the new or modified terms are (continuing to drop *int* subscripts),

$$J_{EE} = - a_b - r - f_N N^* - e_{IN} f_{IN} I^* N^* < 0 \quad \text{(A29.a)}$$

$$J_{IE} = (a_b + e_{IN} f_{IN} E^* N^*) / e_I \quad \text{(A29.b)}$$

$$J_{II} = (e_{IN} f_{IN} E^* N^*) / e_I - m_I \quad \text{(A29.c)}$$

hence, the energy self-limitation ($J_{EE}$) has an additional negative term from baseline allocation (-$a_b$); the effect of energy on immune cells ($J_{IE}$) adds $a_b / e_I$; and the intraspecific effect of immune cells ($J_{II}$) is no longer zero (eq. A29.a-c, respectively). It turns that out that $J_{II}$ is always negative (as far as we can discern) because $E_{int}^*\, N_{int}^* < EN_I^*$ (from. eq. A23) – the stimulatory product of energy and parasites sits below the minimum needed by the parasite without baseline allocation (i.e., when $a_b = 0$). Hence level 1 feedback and level 2 feedback,

$$F_1 = J_{EE} + J_{II} < 0 \qquad \text{(A30.a)}$$

$$F_2 = J_{EN} J_{NE} + J_{IE} J_{EI} + J_{IN} J_{NI} - J_{EE} J_{II} < 0 \qquad \text{(A30.b)}$$

are both always negative. The $F_1 < 0$ result then ensures that $F_2$ is always negative (just based on signs alone). As described in the text, the sign of $F_3$ is difficult to prove, but for a wide range of parameter values yielding a feasible interior, it is only negative. (We ran a hypercube search, with a range of 0.1 – 500 units for each parameter across all combinations; the two interior equilibria were never both feasible).

(D) The stabilizing effect of baseline allocation to the PIE models

As indicated in the main text, the baseline allocation of energy to the immune system ($a_b > 0$) is highly stabilizing to dynamics of energy, *E*, parasite, *N*, and immune cells, *I* (Fig. A2.A). In fact just a bit of baseline allocation can shift dynamics from (possibly complex) cycles (e.g., Fig. A2.C) to stable dynamics (e.g., Fig. A2.D). Also, increasing $a_b$ increases density of immune cells and host energy while depressing parasite density (Fig. A2.B).

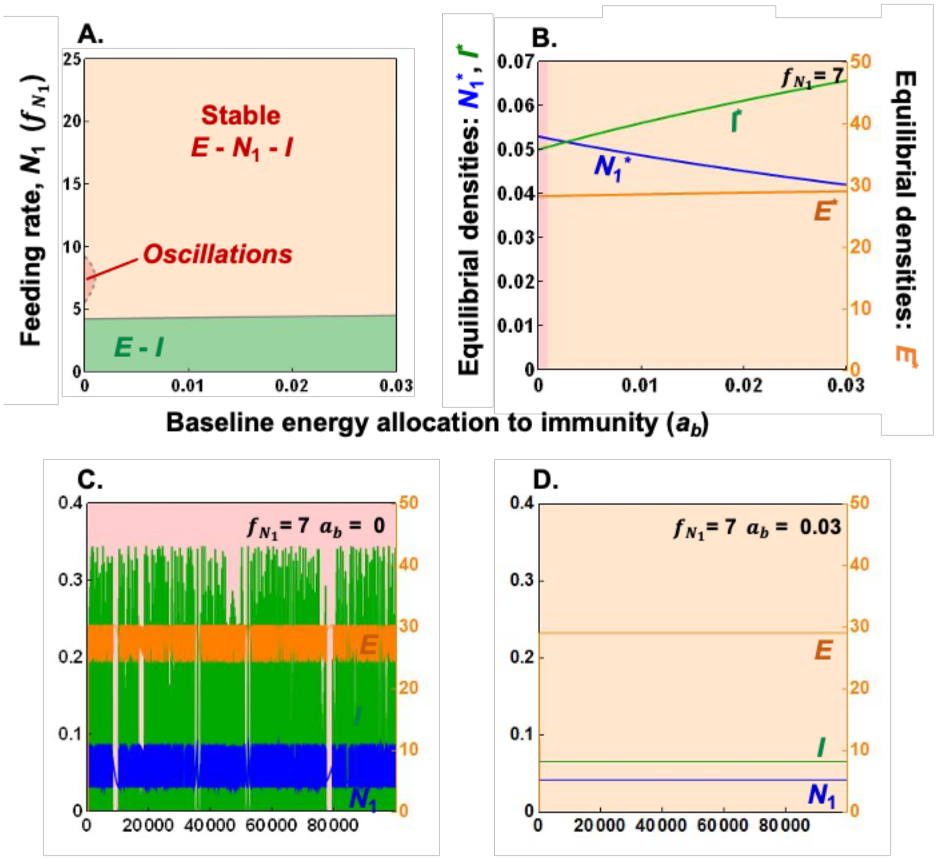


**Figure A2:** *Increasing constitutive immunity eliminates oscillations and lowers parasite burden (density) in PIE models:* (A) Bifurcation diagrams in parameter space of baseline energy allocation to immunity ($a_b$) – feeding rate of parasite ($f_{N_1}$) capture dynamics of PIE ($a_b$ = 0) and PIEc ($a_b$ > 0) for similar parameter values in Fig. 3 (with high resource supply point for hosts [$S$ = 500]). Oscillations (dark orange region) are enveloped within stable coexistence (light orange) regions. (B) *Equilibrial densities along $a_b$* [at $f_{N_1}$ = 7]. Increasing $a_b$ increases density of immune cells, $I^*$ (green) and host energy, $E^*$ (orange) while reducing parasite densities, $N_1^*$ (blue). The densities of *I, E, $N_1$* over time show [C] ($a_b$ = 0) complex oscillatory, or [D] ($a_b$ = 0.03) stable dynamics (see text for details and Table 1 for default parameters).

**Section 3: Analysis of the four dimension models (KP, 2PIE, and 2PIEc)**

In this section, we detail models of mixed apparent and resource competition in the food web (keystone predation, KP) and within a host (2PIE and 2PIEc; Figs. 5,6; Table 2).

*System of equations (see also Table 2)*

Equations for these four dimensional models largely follow their 3D cousins. In KP, the predator ($P$) feeds on the prey ($N_i$) at a rate $f_{PN_i}$ and conversion efficiency $e_{PN_i}$ while experiencing background mortality rate $m_p$ (Eq. 3A). The two prey compete for resources with feeding rate $f_{N_i}$ and conversion rate $e_{PN_i}$; they both have equal background death rate $m_N$ (Eq. 3B). The chemostat-like resource has supply point $S$ and dilution rate $a$; both prey have identical stoichiometric conversion, $Q$ (Eq. 3C). The 2PIE models just add a second parasite ($N_i$) to the PIE models, where the parasite specific parameters are denoted for parasite $i$ (Eq. 4A-C).

(A) Keystone predation (KP)

The keystone predation model provides the food web analogy to the model of within-host competition among parasites. It envisions two competitors, $N_1$ and $N_2$, who share a resource, $R$, but also a predator, $P$. Hence, they engage in both resource and apparent competition, respectively. As described below, we envision a tradeoff, such that $N_1$ is the superior resource competitor without predation but is more susceptible to predation than $N_2$.

*The Jacobian matrix, feedback loops, and key ratios*

The Jacobian matrix for the keystone predation model ($\mathbf{J_{KP}}$), evaluated when at a feasible interior equilibrium (but *int* subscripts dropped for easier reading here) and in order of $R$, $N_1$, $N_2$, and $P$, is:

$$\mathbf{J_{KP}} = \begin{bmatrix} F_1 & -f_{N_1} QR^* & -f_{N_2} QR^* & 0 \\ e_{N_1R} f_{N_1} N_1^* & 0 & 0 & -f_{PN_1} N_1^* \\ e_{N_2R} f_{N_2} N_2^* & 0 & 0 & -f_{PN_2} N_2^* \\ 0 & e_{PN_1} f_{PN_1} P^* & e_{PN_2} f_{PN_2} P^* & 0 \end{bmatrix} \quad \text{(A31)}$$

where feedback at levels 1-3,

$$F_1 = J_{RR} = -a - (f_{N_1} N_1^* + f_{N_2} N_2^*)Q < 0 \quad \text{(A32.a)}$$

$$F_2 = (J_{N_1R} J_{RN_1} + J_{N_2R} J_{RN_2}) + (J_{N_1P} J_{PN_1} + J_{N_2P} J_{PN_2}) < 0 \quad \text{(A32.c)}$$

$$F_3 = J_{RR} \, (J_{N_1P} J_{PN_1} + J_{N_2P} J_{PN_2}) < 0 \quad \text{(A32.c)}$$

are all always negative. By signs alone, the level 2 feedback is always negative because it is the sum of four binary consumer resource loops (consumption of resources by both competitors [first set of terms] and predation upon them [second set of terms]). Similarly, we can determine by sign alone that level 3 feedback is always negative (since it is a product of self-limitation of the resource, $J_{RR}$, times the sum of two predation loops, where each $N_j$ is eaten by $P$; eq. A32.c).

Hence, stability of the interior equilibrium (determining coexistence vs. alternative stable states) depends on the sign of level 4 feedback, $F_4$. As described in the main text, $F_4$ is the sum of two positive (destabilizing) and two negative (stabilizing) loops (Fig. 5). Furthermore, $F_4$ also becomes proportional to:

$$\underbrace{\left(\frac{J_{PN_1}}{J_{RN_1}} - \frac{J_{PN_2}}{J_{RN_2}}\right)}_{effects\ on\ (-\varepsilon_i)} x \underbrace{\left(\frac{J_{N_1P}}{J_{N_1R}} - \frac{J_{N_2P}}{J_{N_2P}}\right)}_{affected\ by\ (-\alpha_i)} \quad \text{(A33)}$$

where we define the *effects on* ratio for each $N_j$, $\varepsilon_j = -\frac{J_{PN_j}}{J_{RN_j}}$, as the ratio of the effects of $N_j$ on its predator ($J_{PN_j}$) vs. its resource ($J_{RN_j}$). (The minus sign keeps the quantity positive, bending the mind less in calculations). These $\varepsilon_j$ are impact vectors *sensu* Leibold (1996). Additionally, we define the *affects on* ratio for each $N_j$, $\alpha_j = -\frac{J_{N_jP}}{J_{N_jR}}$, as the ratio of the effects of the predator and resource on the competitor ($J_{N_jP}$ and $J_{N_jR}$), respectively, with again a minus sign convention to aid the work below. These ratios can be written as:

$$\underbrace{\frac{P^*}{QR^*}\left(\frac{e_{PN_1}f_{PN_1}}{f_{N_1}} - \frac{e_{PN_2}f_{PN_2}}{f_{N_2}}\right)}_{effects\ on\ (\varepsilon_1-\varepsilon_2)} x \underbrace{\left(\frac{f_{PN_1}}{e_{N_1}f_{N_1}} - \frac{f_{PN_2}}{e_{N_2}f_{N_2}}\right)}_{affected\ by\ (\alpha_1-\alpha_2)} . \quad \text{(A34)}$$

We refer to the difference in those *effects on* ratios, $\varepsilon_1 - \varepsilon_2$, and the *affected by* ratios, $\alpha_1 - \alpha_2$, below, after defining some key quantities. As will be shown, the difference in *affected by* ratios guarantees a key tradeoff structure while the difference in *effects on* ratio governs switches in competitive hierarchies (or not). Both provide core components of stability analysis of the interior equilibrium.

*Building the bifurcation diagram*

In the KP model, each competitor *j* has a minimal resource requirement, $R_j^*$. Since (for simplicity) we assume each competitor has the same background mortality, $a+m$, the minimal resource requirement is:

$$R_j^* = \frac{a+m}{e_{N_jR}\, f_{N_j}} \quad \text{(A35)}$$

the ratio of death rate to gains (per resource birth rate; equ. 1). When $S > R_j^*$, competitor $N_j$ can invade the resource-only (boundary) system (crossing the border between the *R* and *R*-$N_j$ in the *S*-$f_{N_1}$ bifurcation diagram: fig. 6A). Once it successfully invades, it reaches equilibrial density:

$$N_{j,b}^* = \frac{a\left(S - R_j^*\right)}{f_{N_j}\, QR{*}_j} \quad \text{(A36)}$$

which is resource production divided by consumption of resources per competitor (eq. A36). Furthermore, $N_1$ outcompetes $N_2$ when it has higher per resource birth rate, $e_{N_1R}\, f_{N_1} > e_{N_2R}\, f_{N_2}$

– that trait arrangement guarantees $N_1$ has the lowest minimal resource requirement, $R_1^* < R_2^*$. (This condition defines the border between the $R$-$N_1$ and $R$-$N_2$ regions [Fig. 6A]).

To build a food chain, the predator can invade this $R_j^*$-$N_{j,b}^*$ boundary system when its minimal prey requirement, $N_j^*$, is met. That minimum is:

$$N_{j,P}^* = \frac{a+m_P}{e_{PN_j} f_{PN_j}} \quad \text{(A37)}$$

which is the ratio of death rate of the predator to its birth rate per prey (eq. A37). Note higher vulnerability to predation (e.g., $f_{PN_1} > f_{PN_2}$) leads to smaller minimal prey requirements (e.g., $N_{1,p}^* < N_{1,p}^*$) assuming equal $e_{PNj}$. When this minimum is met (i.e., the $R$-$N_j$ to $R$-$N_j$-$P$ borders are crossed in Fig. 6A), density of resources and predators both increase with the supply point, $S$:

$$R_{j,P}^* = \frac{aS}{a + f_{N_j} N_{j,P}^* Q} \quad \text{(A38.A)}$$

$$P_j^* = \frac{e_{N_j R} f_{N_j}}{f_{PN_j}} \left(R_{j,P}^* - R_j^*\right) \quad \text{(A38.B)}$$

where resources are set at the ratio of supply to losses from dilution and total consumption (equ. A38.A). All else equal, more vulnerable prey compete less well with predators (i.e., they have higher $R_{j,P}^*$ when they have higher $N_{j,p}^*$). Predator density – and the competitor's $P_j^*$ - increases with the difference between competitive ability of $N_j$ with and without predators, $R_{j,P}^*$ - $R_j^*$. This $R_{j,P}^*$-$N_{j,p}^*$-$P^*$ food chain becomes feasible (i.e., the predator invades) with higher supply point,

$$aS > aR_j^* + f_{N_j} R_j^* N_{j,P}^* Q \quad \text{(A39)}$$

which requires that the rate of resource supply, $aS$, exceeds the loss of resource set at the minimal requirement of the competitor without predation, $R_j^*$, plus loss due to consumption by competitors held at the predator's minimum requirement ($N_{j,P}^*$).

The interior equilibrium ($R$-$N_1$-$N_2$-$P$) allows for coexistence or priority effects. When feasible, density of the resource, predator, two competitors at the interior (*int*) become:

$$R^*_{int} = \frac{(f_{PN1} - f_{PN2})}{f_{PN_1}R^*_1 - f_{PN_2}R^*_2} = \frac{(a+m_N)(f_{PN_1} - f_{PN_2})}{\alpha_1 - \alpha_2} \quad \text{(A40.A)}$$

$$P^*_{int} = \frac{(R^*_2 - R^*_1)}{f_{PN_1}R^*_1 - f_{PN_2}R^*_2} = \frac{(R^*_2 - R^*_1)}{(f_{PN_1} - f_{PN_2})} R^*_{int} \quad \text{(A40.B)}$$

$$N^*_{1,int} = \frac{N^*_{1,P}}{Q}\left(\frac{f_{N2} f_{N_2}}{f_{N2}N^*_{2,P} - f_{N_1}N^*_{1,P}}\right)\left(a + f_{N_2}N^*_{2,P} - \frac{aS}{R^*_{int}}\right) \quad \text{(A40.C)}$$

$$N^*_{2,int} = \frac{N^*_{2,P}}{Q}\left(\frac{f_{N_1} f_{N_2}}{f_{N_2}N^*_{2,P} - f_{N_1}N^*_{1,P}}\right)\left(-a - f_{N_1}N^*_{1,P} + \frac{aS}{R^*_{int}}\right) \quad \text{(A40.D)}$$

For feasibility, specific assumptions are needed. Let us assume that $N_2$ is the more defended competitor, i.e., predators have higher attack rate on $N_1$ than $N_2$, then $f_{PN_1} > f_{PN_2}$. Meanwhile, assume $N_1$ is the superior competitor without predators ($R_1^* < R_2^*$, so $e_{N_1R}\, f_{N_1} > e_{N_2R}\, f_{N_2}$). If that tradeoff holds, then positive resource and predator densities (equ. A40.A,B) require that the ranking of *affected by* ratios must be $\alpha_1 > \alpha_2$. This arrangement ensures that $f_{PN_1}\, R_1^* > f_{PN_1}\, R_2^*$, which in turn requires a strong enough tradeoff in vulnerability vs. competitive ability ($f_{PN_1}$ / $f_{PN_2} > R_2^* / R_1^*$). (The boundary $f_{PN_1}\, R_1^* = f_{PN_2}\, R_2^*$ forms an upper asymptotic limit to the alternative stable state region in $S$-$f_{N_1}$ space of Fig. 6.A, but it sits beyond the range visualized).

The two longer expressions for the interior densities place another bound on feasibility. The first expression in parentheses is proportional to the difference of *effects on* ratios: if $\varepsilon_1 > \varepsilon_2$, then it is positive (because $\varepsilon_1 > \varepsilon_2$ ensures that $f_{N_2}N^*_{2,P} > f_{N_1}N^*_{1,P}$). The second terms in parentheses mark two transcritical thresholds, $T_1$ and $T_2$:

$$T_1\!: S = \frac{R^*_{int}}{a}\left(a + f_{N_2}N^*_{2,P}Q\right), \text{ occurring at } R_{int}^* = R_{2,P}^* \;\&\; P_{int}^* = P_2^* \quad \text{(A41.A)}$$

$$T_2\!: S = \frac{R^*_{int}}{a}\left(a + f_{N_1}N^*_{1,P}Q\right), \text{ occurring at } R_{int}^* = R_{1,P}^* \;\&\; P_{int}^* = P_1^* \quad \text{(A41.B)}$$

These expressions (eq. 41) are also the invasion thresholds for $N_2$ into the $R$-$N_1$-$P$ chain and $N_1$ into the $R$-$N_2$-$P$ chain respectively. If there is symmetry in *affected by* and *effects on* ratios (i.e., if $\alpha_1 > \alpha_2$ and $\varepsilon_1 > \varepsilon_2$) then coexistence occurs within the range $T_2 < S < T_1$. This arrangement guarantees an ordering of competitive abilities for resources with predators, $R_{j,P}^*$, and resource density that supports the diamond web, $R_{int}^*$: $R_{2,P}^* < R_{int}^* < R_{1,P}^*$. In other words, more resistant $N_2$ must become the superior resource competitor with predation, but $R_{int}^*$ must fall in between the minimal requirement of the two food chains. It also guarantees an ordering of apparent competitive abilities for the diamond web: $P_{2,P}^* < P_{int}^* < P_{1,P}^*$. Hence, the resistant species must be the inferior apparent competitor ($P_{2,P}^* < P_{1,P}^*$), but the coexistence $P_{int}^*$ must sit between. Also, in this scenario, along a gradient of $S$, $N_2$ invades at $T_1$. Just below this threshold, more vulnerable $N_1$ wins through apparent competition (higher $P_j^*$ [eq. A38.B]). Then with higher $S$ yet, $N_1$ is excluded at $T_2$. Here, more resistant $N_2$ wins through resource competition (lower $R_{j,P}^*$ [eq. A38.A]).

Alternative stable states can emerge for different combinations of nutrient supply-trait space (like $S$-$f_{N_1}$ shown in Fig. 6.A). But first, those two transcriticals cross ($T_1 = T_2$; equ. 7) with increasing $f_{N_1}$ when $f_{N_1}N_{1,P}^* = f_{N_2}N_{2,P}^*$ (hence $\varepsilon_1 = \varepsilon_2$). However, increasing $f_{N_1}$ further switches ranking of *effects on* ratios, creating an asymmetry. Now, $\varepsilon_1 < \varepsilon_2$, hence $f_{N_2}N_{2,P}^* < f_{N_1}N_{1,P}^*$ . Once this switch happens, alternative stable arises within the range $T_1 < S < T_2$. (Note that a certain tradeoff strength is still needed, $f_{PN_1} / f_{PN_2} > R_2^* / R_1^*$, for feasibility, so $\alpha_1 > \alpha_2$). Now, alternative stable states arise when $R_{int}^*$ falls within the range $R_{2,P}^* > R_{int}^* > R_{1,P}^*$. Notice how resource competitive ordering switched: $N_1$ is the superior competitor with and without predation. That shift is paired with a now-reversed ordering of apparent competitive abilities: $P_{2,P}^* > P_{int}^* > P_{1,P}^*$. Hence, now the more resistant species is best at apparent competition ($P_{2,P}^* > P_{1,P}^*$) while $P_{int}^*$ falls between them. Also, at just lower nutrient supply ($S < T_1$), $N_1$ wins via resource competition; at just higher supply ($S > T_2$), $N_2$ wins by apparent competition.

*Summary:* Therefore, stability of the interior equilibrium, feedback loops, competitive hierarchies, and assembly rules all share deep connections in this model of resource and apparent competition (with two niche dimensions). The sum of two positive and two negative level 4 feedback loops (Fig. 5) is proportional to the difference in two key ratios. The difference in *affected by* ratio ensures that a strong enough tradeoff exists in resource competition ($R_j^*$) and resistance to predation ($f_{PN_j}$) between competitors. Here, since $N_1$ had lower minimal resource requirement ($R_1^* < R_2^*$) but was less resistant ($f_{PN_1} > f_{PN_2}$), a feasible interior required that $N_1$ was more affected by predators – relative to resources – than $N_2$ (so $\alpha_1 > \alpha_2$). The difference in *effects on* ratio then proved more restrictive; it governs whether or not there is symmetry or asymmetry with it and the *affected by* and *effects on* ratios. It also determines if there is a switch in competitive hierarchies with and without predation. Coexistence requires symmetry and a switch. In other words, $\varepsilon_1 > \varepsilon_2$ means that now, for resources, resistant $N_2$ is superior for resources ($R_{1,P}^* > R_{2,P}^*$), but has higher $P^*$ ($P_1^* > P_2^*$). Such an arrangement guarantees that negative feedback dominates ($F_4$) – the stabilizing consumer-resource relationships remain stronger than the destabilizing positive loops. Each competitor exerts negative intraspecific indirect effects ($IE_{N_1} < 0$ & $IE_{N_2} < 0$).

In contrast, alternative stables states arise with asymmetry in the *affected by* vs. *effects on* ratios. Hence, there is no switch in competitive hierarchies. Instead, $\varepsilon_2 > \varepsilon_1$ means that now, $N_1$ competes superiorly for resources always, without predation ($R_1^* < R_2^*$) and with ($R_{1,P}^* < R_{2,P}^*$). However, more vulnerable $N_1$ also is the inferior apparent competitor, as it now has lower $P^*$ ($P_1^* < P_2^*$). This configuration leads to dominance of positive feedback ($F_4 > 0$) because the two destabilizing loops overwhelm the stabilizing effects of binary consumer-resource relationships. Each competitor now exerts indirectly facilitates itself via positive intraspecific indirect effects ($IE_{N_1} > 0$ & $IE_{N_2} > 0$).

Of course, these outcomes (coexistence or alternative stable states) only become possible if the interior equilibrium is feasible. Feasibility, in turn, requires that the diamond web's $R_{\mathrm{int}}^*$ and $P_{int}^*$ fall in between those of its food chain building blocks (i.e., for assembly). Otherwise, one competitor excludes the other no matter what.

(B) Some results from the 2PIE model

The 2PIE model produces many relationships that closely resemble those of its food web analogue (KP). Essentially, the mix of exploitative competition between parasites, $N_j$, for energy ($E$) and apparent competition from immune cells ($I$) follows the same basic logic.

*The Jacobian matrix, feedback loops, and key ratios*

First, the Jacobian matrix for 2PIE, $\mathbf{J_{2PIE}}$, in order of $E$, $N_1$, $N_2$, and $I$ (after dropping '*int*' subscripts) is:

$$\mathbf{J_{2PIE}} = \begin{bmatrix} F_1 & -f_{N_1}E^* & -f_{N_2}E^* & 0 \\ \frac{f_{N_1}N_1^*}{e_{N_1E}} & 0 & 0 & -f_{IN_1}N_1^* \\ \frac{f_{N_2}N_2^*}{e_{N_2E}} & 0 & 0 & -f_{IN_2}N_2^* \\ (\sum e_{IN_j}f_{IN_j}N_j^*)E^* & e_{IN_1}f_{IN_1}I^*E^* & e_{IN_2}f_{IN_2}I^*E^* & 0 \end{bmatrix} \quad \text{(A42)}$$

where feedback at level 1, $F_1$, is:

$$F_1 = J_{EE} = -r - \Sigma f_j(I)N_j < 0 \quad \text{(A43)}$$

and energy consumption per parasite, directly and indirectly via immune stimulation, is $f_j(I) = f_{N,j} + e_{IN,j}\, f_{IN,j}I$. Level 1 feedback is always negative. Notice from $\mathbf{J_{2PIE}}$ (eq. A42) how now the effect of energy on the immune system, $J_{IE}$, is a positive quantity (i.e., the direct stimulatory effect of contact with parasites). Its equivalent in the KP model, $J_{PR}$, was zero at the interior (because there was no direct connection between the resource and the non-omnivorous predator).

With this added direct effect, level 2 feedback simply sums those negative pairwise consumer-resource like relationships:

$$F_2 = (J_{N_1 I} J_{IN_1} + J_{N_2 I} J_{IN_2}) + (J_{N_1 E} J_{EN_1} + J_{N_2 E} J_{EN_2}) + (J_{IE} J_{EI}) < 0 \quad \text{(A44)}$$

which, from left to right, is the sum of immune interactions with both parasites (first terms in parentheses), consumption of energy by both parasites (second terms), and use of energy by the immune system (i.e., that new loop not present in KP). Level 3 feedback is the sum of the three loops present in PIE, (i) + (ii) + (iii) at level 3 (Fig. 2), but now also summed for both parasites $N_j$ (Fig. 4):

$$F_3 = -\Sigma[\, (J_{N_1 I} J_{IN_j} J_{EE}) + (J_{EN_j} J_{IE} J_{IN_j}) + (J_{IN_j} J_{EI} J_{N_j E})\,] < 0 \quad \text{(A45)}$$

Level 3 feedback is always negative. Hence, level four feedback, $F_4$, determines whether the interior equilibrium enables coexistence ($F_4 < 0$) or is a saddle ($F_4 > 0$), one that creates the potential for alternative stable states between the two $E$-$N_j$-$I$ systems. As above for KP, coexistence requires symmetry between *effects on* and *affected by* ratios (modified from eq. A33), where:

$$\underbrace{I^* \left(\frac{e_{IN_1} f_{IN_1}}{f_1(I)} - \frac{e_{IN_2} f_{IN_2}}{f_2(I)}\right)}_{\text{effects on } (\varepsilon_1 - \varepsilon_2)} x \underbrace{\left(\frac{e_{N_1} f_{IN_1}}{f_{N_1}} - \frac{e_{N_2} f_{IN_2}}{f_{N_2}}\right)}_{\text{affected by } (\alpha_1 - \alpha_2)} \quad \text{(A46)}$$

and these quantities are described and used below.

*Building the bifurcation diagram*

Borrowing from analysis of the PIE model above (section 2B), we define key quantities for parasite *j*. The minimal energy requirements for parasite j to invade a host with no standing immune cells (the *E*-only system in Fig. 6B), $E_j^*$, and the parasite density reached, $N_{j,t}^*$, are:

$$E_j^* = \frac{e_{NE,j} m_N}{f_{N.j}} \quad \text{(A47.a)}$$

$$N^*_{j,t} = \frac{f(S)/E^*_j - r}{f_{IN,j}} \qquad \text{(A47.b)}$$

$$EN^*_{j,t} = \frac{f(S) - rE^*_j}{f_{IN,j}} \qquad \text{(A47.c)}$$

Hence the parasite invades (and $N_{j,t}^* > 0$) when $f(S) > r\, E_j^*$ . This equilibrium produces a product of energy and parasites $EN_{j,t}^*$. Immune cells invade when this product exceeds minimal $EN$ requirement of the immune cells, $EN_{I,j}^*$ (i.e., when $EN_{j,t}^* > EN_{I,j}^*$). With this requirement met, the $E$-$N_j$-$I$ boundary system then sets the density of energy ($E_{j,b}^*$), parasite ($N_{j,b}^*$), and immune cells ($I_{j,b}^*$) for the single parasite system:

$$EN^*_{I,j} = \frac{e_{I,}m_I}{e_{IN,j}f_{IN,j}} \qquad \text{(A48.a)}$$

$$E^*_{j,b} = \frac{e_{NE,j}\left(f(S) - (f_{N,j} - e_{IN,j}m_N)EN^*_{I,j}\right)}{e_{IN,j}f_{N,j}EN^*_{I,j} + e_{NE.j}r} \qquad \text{(A48.b)}$$

$$N^*_{j,b} = EN^*_{I,j}/E^*_{j,b} \qquad \text{(A48.c)}$$

$$I^*_{j,b} = \frac{f_{N,j}\left(f(S) - rE^*_{N,j} - f_{N,j}EN^*_{I.j}\right)}{e_{I,j}f_{N,j}m_I + e_{NE.j}r}\,. \qquad \text{(A48.d)}$$

(There is a condition required for $E_{j,b}^* > 0$ [from equ. A48.b], but this condition proves less restrictive than $EN_{j,t}^* > EN_{I,j}^*$).

All four components of this within-host interaction can maintain positive densities at a feasible interior equilibrium. At this interior (*int*), energy and immune cells reach (respectively):

$$E^*_{int} = \frac{f_{N,1}f_{N,2}E^*_1E^*_2(f_{IN,1} - f_{IN,2})}{f_{IN,1}E^*_1 - f_{IN,2}E^*_2} \qquad \text{(A49.a)}$$

$$I^*_{int} = \frac{m_N(E^*_2 - E^*_1)}{f_{IN,1}E^*_{N,1} - f_{IN,1}E^*_{N,2}} \qquad \text{(A49.b)}$$

which, like KP, requires a certain tradeoff structure. We imagine that parasite $N_1$ is the superior energy competitor without immune cells (so $E_1^* < E_2^*$ [from eq. A47.a]) but is also more vulnerable to immune attack (so $f_{IN,1} > f_{IN,2}$). Given these assumptions, a feasible interior requires a strong enough tradeoff so that $f_{IN,1}\, E_{N,1}{}^{*} > f_{IN,2}\, E_{N,2}{}^{*}$. This condition is assured when the *affected by* ratio leads to $\alpha_1 > \alpha_2$ (from eq. 46). Density of the two parasites at the interior, $N_{j,int}{}^{*}$, becomes:

$$N^*_{1,int} = \frac{EN^*_{I,1}}{E^*_{int}}\left(\frac{rE^*_{int}+f_2(N,I)EN^*_{I,2}-f(S)}{f_2(I)EN^*_{I,2}-f_1(I)EN^*_{I,2}}\right) \qquad \text{(A50.a)}$$

$$N^*_{2,int} = \frac{EN^*_{I,2}}{E^*_{int}}\left(\frac{-rE^*_{int}-f_1(N,I)EN^*_{I,1}+f(S)}{f_2(I)EN^*_{I,2}-f_1(I)EN^*_{I,1}}\right) \qquad \text{(A50.b)}$$

where summed per parasite consumption of energy, $f_j(I) = f_{N,j} + e_{IN,j}\, f_{IN,j}\, I_{int}$, is that directly consumed and that activated as immune cells attack parasites. These expressions place additional bounds on feasibility. The denominator is proportional to the difference of *effects on* ratios: if $\varepsilon_1 > \varepsilon_2$ (from eq. A46), then the denominator it is positive (because $\varepsilon_1 > \varepsilon_2$ ensures that $f_2(I)EN^*_{2,P} > f_1(I)EN^*_{2,P}$). The numerators mark two transcritical thresholds, $T_1$ and $T_2$:

$$T_1\text{: } f(S) = rE^*_{int} + f_2(N,I)EN^*_{I,2} \text{ occurring at } E_{int}{}^{*} = E_{2,b}{}^{*} \;\&\; I_{int}{}^{*} = I_{2,\mathrm{b}}{}^{*} \qquad \text{(A51.a)}$$

$$T_2\text{: } f(S) = rE^*_{int} + f_1(N,I)EN^*_{I,1} \text{ occurring at } E_{int}{}^{*} = E_{1,b}{}^{*} \;\&\; I_{int}{}^{*} = I_{1,\mathrm{b}}{}^{*} \qquad \text{(A51.b)}$$

Hence, a now familiar story arises for resource and apparent competition between the two parasites. Parasite $N_1$, the superior resource competitor without immune cells ($E_1{}^{*} < E_2{}^{*}$), can coexist with $N_2$, the parasite which better resists immune attack ($f_{IN,1} > f_{IN,2}$). With symmetry of *effects on* and *affected by* ratios, $\alpha_1 > \alpha_2$ and $\varepsilon_1 > \varepsilon_2$, the parasites coexist when incoming energy flow, $f(S)$, falls between $T_1 < f(S) < T_2$. In this range, a ranking of competitive ability for energy emerges, $E_{2,b}{}^{*} < E_{int}{}^{*} < E_{1,b}{}^{*}$. Notice how competitive abilities for energy have flipped with immune attack (but the coexistence $E^*$ value falls in between those single-parasite values). Additionally, in this range, $I_{2,b}{}^{*} < I_{int}{}^{*} < I_{1,b}{}^{*}$. This ranking means that the less resistant parasite ($N_1$) is the superior apparent competitor because it can support higher density of immune cells. Also, the coexistence $I^*$ falls in between these values of immune cells supported by singly infected hosts. Hence, symmetry in these key ratios ensures a strong enough tradeoff ($\alpha_1 > \alpha_2$)

and a flip in competitive hierarchy with immune cells ($\varepsilon_1 > \varepsilon_2$) to ensure coexistence within a certain range of energy supply to the host, *f*(*S*). Furthermore, those conditions lead to each species exerting negative intraspecific indirect effects on themselves ($IE_{N_1} < 0$ and $IE_{N_2} < 0$). At low energy supply, $f(S) < T_1$, more vulnerable parasite $N_1$ wins by apparent competition; at high energy supply, $f(S) > T_2$, more resistant $N_2$ wins by resource competition.

In contrast, alternative stable states between the two parasites can arise when $N_1$ has even higher feeding rate on the resource ($f_{N_1}$). This situation can create asymmetry in *effects on* and *affected by* ratios in the range $T_2 < f(S) < T_1$. Within this range, now the tradeoff remains strong enough ($\alpha_1 > \alpha_2$ still) but a shift in competitive hierarchies emerges: now, $\varepsilon_1 < \varepsilon_2$, which means that $f_2(I)EN^*_{I,2} > f_1(I)EN^*_{I,1}$. With this shift, parasite $N_1$ remains the superior energy competitor with or without immune cells; it wins via exploitative competition for energy when *f*(*S*) is below $T_2$ and within the alternative stable states region. More resistant parasite $N_2$, in turn, supports more immune cells and wins by apparent competition, when *f*(*S*) is above $T_1$ but also in the alternative stable states region. In that region, each species exerts positive intraspecific indirect effects on themselves ($IE_{N_1} > 0$ *and* $IE_{N_2} > 0$). Hence, at this interior equilibrium, positive feedback leads to exclusion of one parasite by the other.

<u>(C) Two four species transitions in KP (and 2PIE) models</u>

Both competitors – prey or parasite – can potentially coexist with their resource and enemy. Above, we outline more typical assembly, via invasion of one of the competitor, $N_i$, into the *R*-$N_j$-*P* food chain or its within-host analogue (*E*-$N_j$-*I*). The less typical assembly arises at lower supply point, *S*, but with increasing feeding rate of $N_1$, $f_{N_1}$ for both KP and 2PIE. At low $f_{N_1}$, $N_1$ and $N_2$ can become competitively equivalent without enemies. Remembering that it controls competitive ability of $N_1$, when $f_{N_1}$ sits too low, $N_2$ also outcompetes $N_1$ for nutrients without enemies (i.e., $N_2$ enjoys lower $R^*$ or $E^*$ - hence $R$-$N_2$ and $E$-$N_2$ regions appear in bottom-left corners of the bifurcation diagrams). Yet, here, more resistant $N_2$ cannot support the predator (in KP: Fig. 6G), or the product of $N_2^*$ $E^*$ cannot maintain production of immune cells (in 2PIE: Fig 6H). Then, with higher $f_{N_1}$, $N_1$ and $N_2$ become competitively equivalent. Here, their composition could range from 100% $N_1$ to 100% $N_2$ or some combination in between. Yet, a particular summed density of more of both prey at the resource only boundary, $N_{j,b}^*$ (eq. A36) relative to the minimal prey requirement of the parasite, $N_{j,P}^*$ (eq. A37):

$$\sum_j \left(N_{j,b}^*/N_{j,P}^*\right) > 1 \qquad \text{(A52)}$$

allows the predator to invade (creating the jump from two to four species coexistence [Figs. 6A,G]). Similarly, a sum of energy times parasite density provided by the $E$-$N_j$ system, $EN_{j,t}^*$ (eq. A47.c), relative to minimal $EN$ requirements of the immune system, $EN_{j,I}^*$ (eq. A48.a)

$$\sum_j \left(EN_{j,t}^*/EN_{j,I}^*\right) > 1 \qquad \text{(A53)}$$

enables invasion of immune cells (Fig. 6B,H). That combination creates the jump from 2 to 4 dimensional stability (but notice how $R_1^* = R_2^* = R_{int}^*$ [eqs. A235, A40.A; Fig. 6.G] and $E_1^* = E_2^* = E_{int}^*$ [eqs. A47.a, A49.a; Fig. 6.H] between both 2 and 4 dimensional systems at those borders).

<u>(D) More about stability analysis, inverse Jacobians, and indirect effects in four dimensions</u>

Let's rewrite the Jacobian matrix for keystone predation, $\mathbf{J_{KP}}$, reordered now as $R$, $P$, 1 [for $N_1$] and 2 [for $N_2$]:

$$\mathbf{J_{KP}} = \begin{bmatrix} J_{RR} & 0 & J_{R1} & J_{R2} \\ 0 & 0 & J_{P1} & J_{P2} \\ J_{1R} & J_{1P} & 0 & 0 \\ J_{2R} & J_{2P} & 0 & 0 \end{bmatrix} \text{ with sign structure } \begin{bmatrix} - & 0 & - & - \\ 0 & 0 & + & + \\ + & - & 0 & 0 \\ + & - & 0 & 0 \end{bmatrix} \qquad \text{(A54)}$$

Feedback for this Jacobian matrix at level 3 ($F_3$) and level 4 ($F_4$) are, respectively:

$$F_3 = - J_{RR}\,(J_{P2}\,J_{2P}) - J_{RR}\,(J_{P1}\,J_{1P}) \qquad \text{(A55.a)}$$

$$F_4 = - (J_{1P}\,J_{2R} - J_{2P}\,J_{1R})\,(J_{P1}\,J_{R2} - J_{P2}\,J_{R1}) = - A\,E \qquad \text{(A55.b)}$$

where $A = - (J_{1P}\,J_{2R} - J_{2P}\,J_{1R})$ and contains information in the difference in the *affected by* ratios (and $A > 0$ guarantees the tradeoff considered above). Similarly, $E = - (J_{P1}\,J_{R2} - J_{P2}\,J_{R1})$ contains information in the difference in the *effects on* ratios (so $E > 0$ leads to coexistence [$F_4 < 0$] while $E < 0$ produces alternative stable states [$F_4 > 0$] for feasible interior equilibria). In four dimensions, due to how signs work with matrices, the inverse Jacobian, $\mathbf{J}^{-1}$ (rather than $-\mathbf{J}^{-1}$) contains the matrix of indirect effects. This inverse Jacobian (remembering the new order $R$, $P$, $N_1$, and $N_2$) is:

$$\mathbf{J}^{-1}{}_{\mathbf{KP}} = \begin{bmatrix} 0 & 0 & J_{2P}/A & J_{1P}/A \\ 0 & 0 & -J_{2R}/A & -J_{1R}/A \\ J_{P2}/E & -J_{R2}/E & -J_{RR}(J_{P2}J_{2P})/(AE) & J_{RR}(J_{P2}J_{1P})/(AE) \\ -J_{P2}/E & J_{R1}/E & J_{RR}(J_{P1}J_{2P})/(AE) & -J_{RR}(J_{P1}J_{1P})/(AE) \end{bmatrix}. \qquad \text{(A56)}$$

This matrix has an informative structure. First, notice how intraspecific indirect effects for competitor $N_j$, $IE_{Nj}$, lie on the main diagonal (trace) and is a ratio of feedback at levels 3 and 4 (the two highest); in other words, tr[$\mathbf{J}^{-1}{}_{\mathbf{KP}}$] = - $F_3/F_4 = IE_{N_1} + IE_{N_2}$ (This point applies to the 2PIE and 2PIEc models, also [not shown]). Hence, the sum of intraspecific indirect effects in both three and four dimensions involves ratios of the two highest levels of feedback (see also section 1 above).

Second, we note some observations within this inverse Jacobian matrix (eq. 56). If we call the upper right quadrant the 'indirect *effect of* niche' submatrix (**IEN**), the lower left the 'indirect *affected by* niche (**IAN**)' submatrix, and the lower right the 'indirect competition' submatrix (**IC**)',

$$\mathbf{J}^{-1}{}_{\mathbf{KP}} = \begin{bmatrix} \mathbf{0} & \mathbf{IEN} \\ \mathbf{IAN} & \mathbf{IC} \end{bmatrix} \tag{A57}$$

we see several interesting patterns based on signs alone in coexistence (CO) vs. alternative stable states (AS) cases. For the indirect competition submatrix (**IC**),

$$\mathbf{IC}_{\mathbf{CO}} = \begin{bmatrix} - & + \\ + & - \end{bmatrix} \text{ while } \mathbf{IC}_{\mathbf{AS}} = \begin{bmatrix} + & - \\ - & + \end{bmatrix} \tag{A58}$$

which says that species coexist when they exert negative intraspecific indirect effects on themselves (main diagonal) but positive effects on their competitors (off diagonal). Conversely, they show alternative stable states when they indirectly facilitate themselves but indirectly harm their competitors. Looking at the 'indirect *affected by* niche' submatrix (**IAN**), we see:

$$\mathbf{IAN}_{\mathbf{CO}} = \begin{bmatrix} + & + \\ - & - \end{bmatrix} \text{ while } \mathbf{IAN}_{\mathbf{AS}} = \begin{bmatrix} - & - \\ + & + \end{bmatrix} \tag{A59}$$

which means that, when competitors coexist, resources and predators indirectly benefit $N_1$ but harm $N_2$; these signs flip with alternative stable states. Finally, the 'indirect *effects on* niche' submatrix (**IEN**) has only one sign structure:

$$\mathbf{IEN}_{\mathbf{CO}} = \mathbf{IEN}_{\mathbf{AS}} = \begin{bmatrix} - & + \\ - & + \end{bmatrix} \tag{A60}$$

which says that $N_1$ has (indirect) negative effects on resources and predators while $N_2$ has (indirect) positive effects on them.
